# Haplotype-resolved long-read transcriptomics uncovers allele-specific heat stress induction and a nucleus-targeted chaperone isoform in potato

**DOI:** 10.64898/2026.11.14.766623

**Authors:** Nadja Nolte, Valentina Levak, Tim Godec, Tjaša Lukan, Katja Stare, Kristina Gruden, Marko Petek

## Abstract

Tetraploid potato is highly heterozygous, and its four haplotypes differ in gene content and regulation of gene and isoform expression, yet responses of individual haplotypes to environmental stresses remain poorly characterized. Here, we used long-read RNA sequencing to profile the haplotype-resolved leaf transcriptome of potato cv. Désirée under prolonged heat. Long-read RNA-seq gene counts correlated with short-read RNA-seq at the gene level but assigned far more reads uniquely to individual haplotypes. Under control conditions, allelic imbalance was widespread, whereas after prolonged heat, most alleles responded in the same way across haplotypes. Allele-specific exceptions point to candidate alleles for heat tolerance. We identified a heat-inducible, redox homeostasis-related N²-acetylornithine deacetylase allele and confirmed its allele-specific promoter activity *in planta*. Heat stress induced alternative isoform usage of 390 genes. Among these, we identified a heat stress upregulated heat shock protein 40 chaperone gene isoform that, unlike its alternative splicing isoform, localizes to the nucleus instead of to the cytoplasm. These results show that in a polyploid crop haplotype-resolved long-read transcriptomics can help in identification of alleles for stress tolerance breeding.

## INTRODUCTION

Heat stress presents a major challenge to global food security, increasingly affecting the productivity of staple crops such as tetraploid potato (*Solanum tuberosum*)^1,2^, the fourth most produced food crop worldwide^3^. Potato is adapted to cool climates and is therefore particularly heat-sensitive. Already moderately elevated temperatures impair tuberization and reduce potato yield^1,4,5^. Currently, differential expression analysis in stress response studies is widely performed at the gene level, which fails to reveal two layers of expression regulation: allele-level expression, whereby the individual alleles of a gene could respond differently, and isoform-level expression, whereby alternatively spliced transcripts could change their relative abundance. Resolving gene expression further to allele and isoform level can reveal new mechanisms and identify targets for breeding more resilient cultivars.

In plants, heat activates a conserved network of heat stress transcription factors (HSFs) that induce heat shock proteins (HSPs), ROS-scavenging enzymes and further HSFs^6,7^. Alternative splicing adds a further layer of complexity, modulating HSF function or targeting transcripts for nonsense-mediated decay^8,9^. The potato heat response has so far been characterized mostly at the gene level, and alternative splicing under heat has so far been profiled only after short-term heat treatment^10^. This leaves open the question which differential isoform usages occur in potato under agronomically relevant prolonged heat stress and whether expression differs between the four alleles of heat-responsive genes. Genome sequences of highly inbred varieties, such as the collapsed doubled monoploid (DM) genome in the case of potato, are commonly used as reference for quantification of gene expression^11^, but cannot resolve cultivar-specific genes, structural variation, or individual haplotypes. Haplotype-resolved reference genomes, now becoming available for crop cultivars^12^, capture cultivar and haplotype-level variation and allow all haplotype copies (alleles) of each gene to be quantified individually, enabling allele-specific expression analysis. In a haplotype-resolved assembly, alleles can be identified as syntelogs, i.e., homologous genes located in collinear regions (the same locus) on different haplotypes; we refer to all alleles of one gene as a syntelog group. A key limitation of phased reference genomes, however, is that the high sequence similarity between haplotypes causes a large fraction of reads from commonly used short-read RNA-seq to multimap, meaning they cannot be uniquely assigned to a single haplotype^13^. Long-read RNA-seq overcomes this by sequencing full-length transcripts: reads spanning entire transcripts map accurately to highly similar haplotypes and also allow novel transcripts and genes to be identified and quantified^14^. Correct read assignment is especially important because many stress-responsive genes belong to large families, so paralogs (gene family members at different loci) and alleles are readily confounded unless reads can be assigned to a defined haplotype.

Allelic imbalance, where the alleles of a gene do not contribute equally to its total expression, can reveal the effects on gene expression caused by allelic variation in *cis*-regulatory regions, such as single nucleotide polymorphisms (SNPs) in transcription factor binding sites or larger structural variants (SVs)^15^. For example, in potato, a quantitative trait locus (QTL) for tuber yield at elevated temperature was resolved by QTL mapping to the heat shock protein *HSC70* allele carrying a TA repeat in its promoter. When expressed under its native promoter in potato cv. Désirée, this allele raised *HSC70* expression and heat tolerance^16^. This shows that individual *cis*-regulatory alleles can have substantial phenotypic consequences. Unlike allopolyploids, which combine genomes from different species and often show subgenome expression bias^17,18^, autotetraploid potato lacks such subgenome dominance because all four haplotypes derive from the same species. Nevertheless, local allelic imbalance can still arise, but has so far been studied only in three potato studies: first in cv. Atlantic, against the collapsed DM reference^19^, and the remaining two using the haplotype-resolved cv. Atlantic reference and short-read RNA-seq^20,21^. Long-read RNA-seq has recently been applied for allele-specific expression analysis in hexaploid sugarcane^22^, but high-depth long-read data remain scarce, and it has not yet been used to resolve allele– and isoform-level expression under stress in any crop. Long-read quantification alone does not explain why expression of alleles differs. Thus, to trace allelic expression differences back to *cis*-regulatory variation, allele-specific expression analysis needs to be combined with haplotype-resolved promoter sequence and chromatin accessibility analyses.

Here, we aim to resolve the response of tetraploid potato to prolonged heat stress at the level of individual alleles and isoforms, and to identify the *cis*-regulatory variation that causes alleles of the same gene to be expressed differently. We hypothesize that (i) long-read RNA-seq mapped to a haplotype-resolved reference genome enables reliable allele-specific quantification; (ii) allelic imbalance is widespread and is at least partly explained by sequence and structural variation in allele promoters, and is therefore shared between cultivars carrying the same haplotypes; and (iii) prolonged heat alters the relative expression of alleles and the usage of isoforms, with consequences for the function of the encoded proteins. To test these hypotheses, we combined long-read RNA-seq of Désirée under 7 days of heat with a haplotype-resolved genome, promoter variation, chromatin accessibility and long-read RNA-seq of a second cultivar, Atlantic, and validated selected predictions experimentally.

## RESULTS

### Long-read RNA-seq reliably quantifies global allele-specific gene expression in potato

To study the expression response of potato to prolonged heat stress at the level of alleles and isoforms, we sequenced leaf samples from control and 7-day heat-treated potato Désirée with PacBio Kinnex long-read RNA-seq. To evaluate quantification accuracy, we first compared our long-read RNA-seq to standard Illumina RNA-seq performed on the same samples (Fig. 1a, Supplementary Table S1). Reads per sample ranged from 37.9M to 50.0M for short-read RNA-seq and from 24.9M to 38.2M for long-read RNA-seq. Mapping long-read RNA-seq to the haplotype-resolved Désirée reference genome identified 25,639 novel isoforms and 6,054 novel genes missing from the UniTato DM annotation^23^ (Supplementary Fig. S1a). 92% of these novel genes were already annotated in Désirée-specific genome annotation^24^, suggesting they represent cultivar-specific genes absent from the DM reference. Long-read data also corrected the intron–exon structure of several mis-annotated genes (Supplementary Fig. S1b). Next, using only single-isoform genes, we assessed how read length and the reference genome affect the number of reads mapped uniquely to one locus. With both technologies, most genes in the collapsed DM reference had a unique mapping rate above 90%, i.e. more than 90% of their reads mapped uniquely (Fig. 1b). Against the haplotype-resolved Désirée genome, however, only 7% of alleles had a unique mapping rate above 90% with short-read RNA-seq, and most alleles had more than 90% multimapping reads, as reads could not be distinguished between highly similar alleles. In contrast, with long-read RNA-seq, 44% of alleles still had a unique mapping rate above 90% on the haplotype-resolved reference (Fig. 1b).

**Fig. 1:**
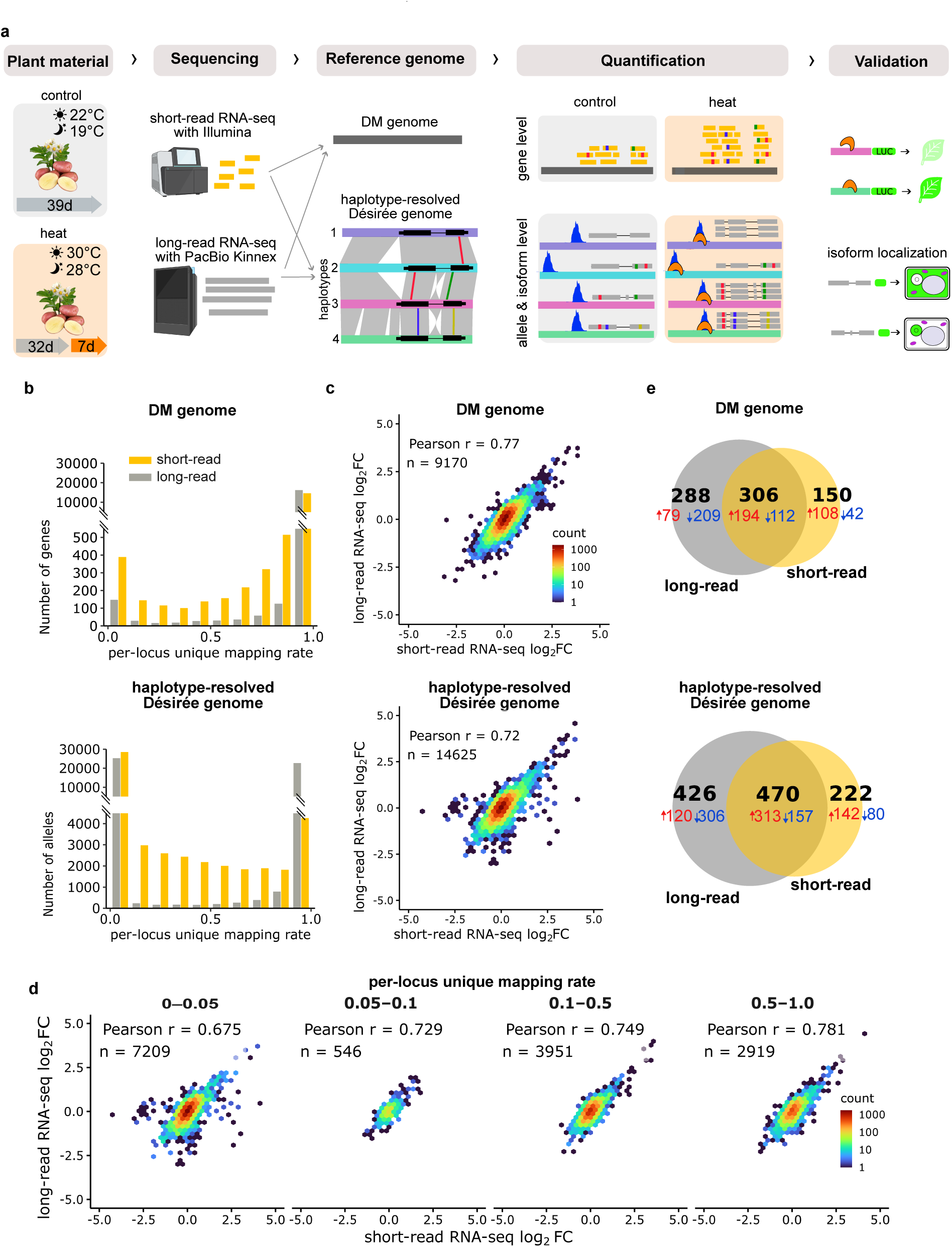
Experimental design and comparison of long-read and short-read RNA-seq in potato Désirée using the collapsed doubled monoploid (DM) and a haplotype-resolved reference. **(a)** RNA samples from potato Désirée under control and 7-day heat-treated conditions were sequenced with Illumina short-read RNA-seq and PacBio Kinnex long-read RNA-seq, and reads were mapped to the collapsed DM reference genome and to the haplotype-resolved Désirée reference genome. Expression is quantified at the gene (top right) or allele and isoform level (bottom right) and can be interpreted with chromatin accessibility data (ATAC-seq coverage in blue) and transcription factor binding (shown in falcate shape). **(b)** Barplots of per-locus unique mapping rate of single-isoform genes for long-(grey) and short-read RNA-seq (yellow), mapped to the collapsed (top) and haplotype-resolved reference (bottom). Higher rates indicate that more reads were assigned uniquely to a gene. **(c)** Correlation of heat/control log2 fold changes (log_2_FC) between short– and long-read RNA-seq, quantified using collapsed (DM, top) and haplotype-resolved (Désirée, bottom) reference genome. In all scatterplots (c, d), only genes detected by both technologies were included; n = number of single-isoform genes. **(d)** Correlation of log2FC between long– and short-read RNA-seq mapped to the haplotype-resolved reference for genes grouped by short-read RNA-seq per-locus unique mapping rate into four groups. **(e)** Venn diagrams of genes differentially expressed between heat and control conditions, detected exclusively by long-read RNA-seq, exclusively by short-read RNA-seq, or by both technologies mapping to the DM reference genome (top) or haplotype-resolved Désirée genome (bottom).

Log_2_FC correlation between technologies was influenced by transcript length: genes shorter than 750 nt showed poor inter-technology correlation, while genes longer than 750 nt showed comparably strong agreement (Supplementary Fig. S2). On the haplotype-resolved Désirée reference genome, correlation between technologies was lower than when using a collapsed genome as reference (Fig. 1c). This drop can be explained by mapping behavior: per-locus unique mapping rate strongly influenced inter-technology correlation, with lower unique mapping rates of short-read RNA-seq corresponding to lower correlation (Fig. 1d). Because the four Désirée haplotypes are highly similar, short reads often multimapped across haplotypes, whereas full-length long reads remained distinguishable. Thus, the lower correlation on the phased reference reflects haplotype-level multimapping of short reads, a limitation that long-read RNA-seq overcomes.

Following this stringent quality control, we identified heat-responsive genes by differential expression analysis and asked whether the sequencing technology affected which genes were detected. No genes were regulated in opposite directions between technologies (Fig. 1e; Supplementary Tables S2–S5), and the functional ontology terms enriched among genes differentially expressed under prolonged heat were largely shared across technologies and reference genomes (Supplementary Table S6). Thus, the sequencing technology only affected the number of genes detected as differentially expressed, with long-read RNA-seq detecting more.

### Allelic imbalance is widespread under control conditions

To evaluate the effect of different haplotypes on gene expression, we compared expression levels of alleles across haplotypes. In the Désirée assembly, 61% of protein-coding genes were found in syntelogous relationships on at least two haplotypes without copy number variation, with 41% present once in all four haplotypes (Fig. 2a, Supplementary Table S7). Syntelog groups present once on each of the four haplotypes were further filtered to keep only sufficiently expressed groups (see Methods) and to remove groups in which two or more alleles have identical sequences. Reads from identical alleles cannot be assigned to a single allele, so allelic imbalance cannot be estimated for them. The 1,915 filtered syntelog groups were tested for allelic imbalance and 803 showed imbalanced expression. No expression bias toward a particular haplotype was observed for any chromosome (Fig. 2b, Supplementary Table S8), which is in line with what is known for autopolyploids. Allele-specific expression results were largely concordant between long– and short-read RNA-seq; at the same false discovery rate (FDR) both detected a similar number of imbalanced syntelog groups. However, allelic ratio differences were systematically smaller with short reads, so more syntelog groups passed the effect-size threshold with long reads (Supplementary Fig. S3, Supplementary Note S1, Supplementary Table S9).

**Fig. 2:**
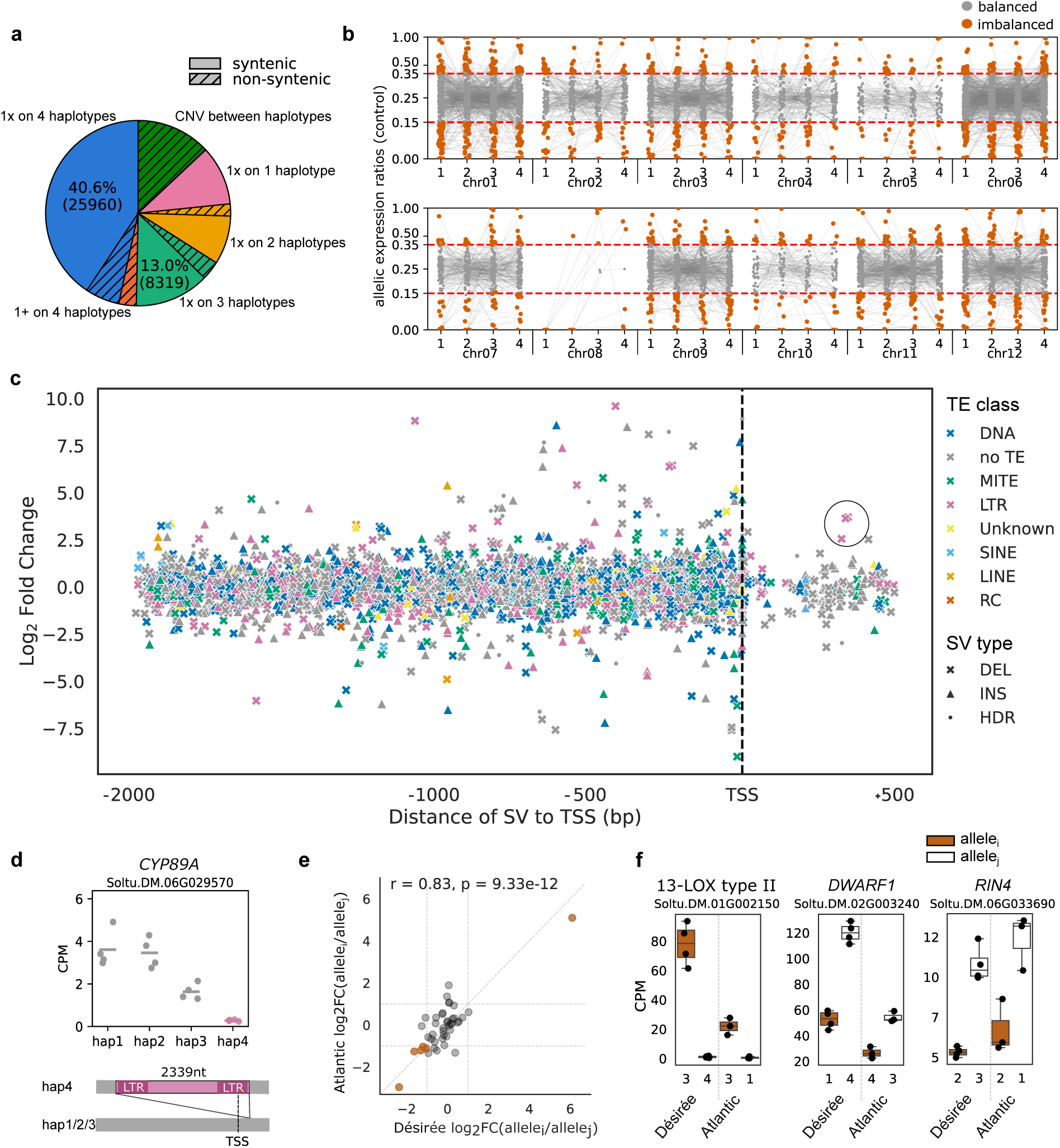
Allelic expression in potato leaves grown under control conditions. **(a)** Pie chart showing the number of genes in the haplotype-resolved Désirée gene annotation classified by their presence/absence and copy number per haplotype in a syntelogous or non-syntelogous relationship (hatched). 1+, the same number of copies (more than one) on each haplotype; CNV, copy number variation, i.e. different numbers of gene copies on different haplotypes. **(b)** Allelic expression ratios of syntelog groups with one allele on each of the four haplotypes with balanced (grey) and imbalanced (orange) expression under control conditions across all potato chromosomes. Red line represents the cutoff for allelic expression ratios defined as imbalanced. **(c)** Difference in expression of alleles (allelic log_2_FC) plotted against the distance of promoter structural variants (SVs) to the transcription start site (TSS, dotted line), type of SV is indicated by color (green: miniature inverted-repeat transposable element (MITE), dark blue: DNA transposon, pink: long terminal repeat (LTR) retrotransposon, orange: long interspersed nuclear element (LINE), grey: not identified as a transposable element (TE), light blue: short interspersed nuclear element (SINE), red: rolling-circle transposon (RC; Helitron), yellow: unknown TE class) and shape (x: deletion, Δ: insertion, °: highly divergent region (HDR)). Circled dots: deletion in promoters of gene shown in panel (d). **(d)** Example of a gene with allelic imbalance – *CYP89A* (Soltu.DM.06G029570). Barplot of allele-specific expression and illustration showing insertion of long terminal repeat (LTR) in the promoter of haplotype 4 (hap4). **(e)** Correlation of allelic log_2_FC between Désirée and Atlantic for genes in which two alleles, i and j, have promoters that are identical between the two cultivars (the promoter of allele i is identical in Désirée and Atlantic, and the same applies to allele j); allelic log_2_FC = log2(CPM of allele i / CPM of allele j), calculated in each cultivar under control conditions; the six genes with allelic imbalance conserved in the same direction in both cultivars are highlighted in orange; r, Pearson correlation coefficient; p, p-value of the correlation. **(f)** CPMs for Atlantic and Désirée of three genes highlighted in (e): *13-LOX type II* (Soltu.DM.01G002150), *DWARF1/DIMINUTO* (Soltu.DM.02G003240, *DIM*), RPM1-interacting protein *RIN4* (Soltu.DM.06G033690). Alleles sharing the same promoter sequence across cultivars are plotted in the same color (allele i, brown; allele j, white). CPM – counts per million.

### Promoter structural variants are frequent but can rarely be linked to allelic imbalance under control conditions

To link differences in promoter sequences to differences in allelic expression, promoter sequences of allele pairs were annotated for larger structural variants (SVs; ≥ 50 bp) and smaller variants (SNPs and indels; < 50 bp). We identified 9,155 promoter pairs with divergent sequences, while 123 promoter pairs were identical (Supplementary Fig. S4a). Of these, 4,340 contained no SVs, 3,318 contained exactly one SV with a median length inside the promoter of 347 bp, and 1,497 contained more than one SV (Supplementary Fig. S4b, Supplementary Table S10). SVs were then classified against a Désirée transposable element (TE) library (Fig. 2c). Most SVs were derived from DNA transposons, followed by long terminal repeat (LTR) retrotransposons, while 42% did not match any TE class (Supplementary Fig. S5). Manual inspection of selected imbalanced allele pairs revealed cases in which an SV in the proximal promoter likely explains the imbalance. For example, the cytochrome P450 gene *CYP89A* (Soltu.DM.06G029570) showed lower expression of haplotype 4 than the other haplotypes (Fig. 2d). Promoter comparison revealed a 2,339 bp insertion 339 bp downstream of the transcription start site (TSS) on haplotype 4, classified as an LTR (Fig. 2d). Across all allele pairs, imbalanced promoter pairs had slightly more SVs than balanced pairs (mean 0.92 vs. 0.80, p = 1.97 × 10⁻⁷), and their SVs lay closer to the TSS (mean 612 vs. 676 bp, p = 3.57 × 10⁻⁵). Among promoter pairs without SVs, imbalanced pairs also had slightly more SNPs and indels than balanced pairs (mean 55.1 vs. 50.7, p = 2.69 × 10⁻³), and their first SNP or indel lay closer to the TSS (mean 72 vs. 85 bp, p = 0.018; Supplementary Fig. S4c). Thus, although promoter sequences of alleles diverge widely in potato, these differences were only weakly associated with allelic imbalance under control conditions, and only in a few cases could a specific variant be linked to it.

### Alleles with identical promoters show correlated imbalance in Désirée and Atlantic

To assess whether allelic expression patterns are maintained across cultivars, allelic expression was compared between cvs. Désirée and Atlantic, reusing publicly available leaf Oxford Nanopore Technologies (ONT) data from cv. Atlantic grown under comparable non-stressed conditions^25^. Out of 939 genes tested pairwise, also testing alleles present not on all four haplotypes, for allelic imbalance in Atlantic and 2,990 in Désirée, 185 genes were shared. Out of these, 43 had an identical pair of promoter sequences in both cultivars, allowing us to compare the expression ratios of alleles with identical promoters in different potato cultivars. The high correlation of log_2_FC for these allele pairs between the two cultivars suggests that alleles with identical promoters exhibit similar relative expression patterns across potato cultivars (Pearson’s r = 0.83, p-value = 9.33 × 10⁻¹²) (Fig. 2e, Supplementary Table S11). Six genes showed significant allelic imbalance in the same direction in both cultivars. Among these were lipoxygenase (*13-LOX type II*) (Soltu.DM.01G002150), cell elongation protein *DWARF1/DIMINUTO* (Soltu.DM.02G003240, *DIM*) and RPM1-interacting protein *RIN4* (Soltu.DM.06G033690) (Fig. 2f). In 4 of the 6 allele pairs with identical promoters whose log_2_FC diverged strongly between cultivars (log_2_FC >1), at least one allele of the gene lacked a gene model in one of the two reference annotations, so its reads were likely counted toward the annotated allele, which may bias the expression ratio (Supplementary Table S12).

### Prolonged heat alters the expression of 1,258 alleles, and a distal promoter region drives stronger heat induction of a NAOD allele

We identified 1,258 alleles, belonging to 734 genes, with changed expression when plants were exposed to prolonged heat stress (Fig. 3a). Functional enrichment analysis of differentially expressed genes identified secondary metabolism (phenolics, terpenoids), cell wall lignin biosynthesis, cytokinin degradation, protease inhibitors and proteolysis, pathogen defense, phytosterol metabolism, NRT1/PTR anion transport, and the enzyme classes oxidoreductases, glycosyltransferases and hydrolases as functionally enriched (Supplementary Table S6).

**Fig. 3:**
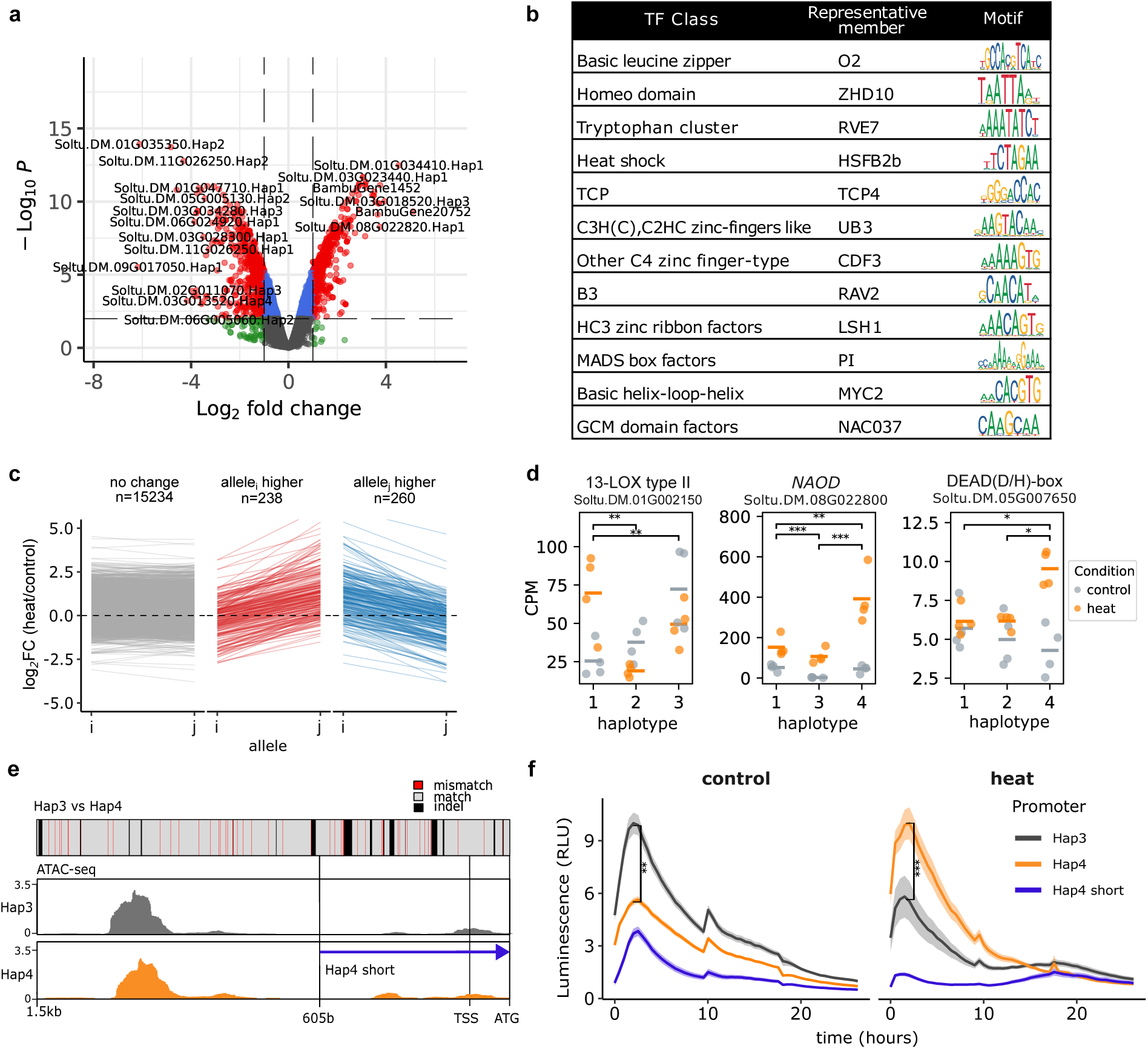
Changes in expression during heat at allele level. **(a)** Volcano plot of differential expression analysis at the allele level, comparing heat versus control conditions. Alleles with |log_2_FC| > 3 are labeled. **(b)** Enriched motifs (Q ≤ 0.05) in promoters of strongly upregulated alleles during heat treatment (log_2_FC > 2). **(c)** Pairwise comparison of the heat response of two alleles of the same gene. Each allele pair (i, j) is shown once, with allele i on the lower-numbered haplotype (hap1–hap3) and allele j on the higher-numbered haplotype (hap2–hap4); log_2_FC is heat/control for each allele, and Δlog_2_FC = log_2_FC(i) − log_2_FC(j). Panels are split by type of expression change. In grey: alleles with a similar response (|Δlog_2_FC| ≤ 1 or FDR ≥ 0.05). In red and blue: alleles with a significantly different response to heat (|Δlog_2_FC| > 1, FDR < 0.05)**. (d)** Example genes with differential allelic expression in heat: a lipoxygenase (13-LOX type II) (Soltu.DM.01G002150), an N²-acetylornithine deacetylase (*NAOD*) (Soltu.DM.08G022800), and a DEA(D/H)-box RNA helicase (Soltu.DM.05G007650). Scatterplot of CPMs per allele in control (grey) and heat (orange); the x-axis indicates the haplotype on which each allele is located. Lines indicate mean of biological replicates. **(e)** Pairwise alignment of promoter sequences for N²-acetylornithine deacetylase (*NAOD*) (Soltu.DM.08G022800) of haplotype 3 (grey), haplotype 4 (orange) and a shorter (blue; 605 bp long) variant of haplotype 4. Identical promoter regions in grey, mismatches in red and small insertions/deletions in black. ATAC-seq coverage of heat-treated replicates for the promoters of haplotypes 3 and 4. **(f)** Luciferase assays under control and heat conditions for N²-acetylornithine deacetylase (*NAOD*) (Soltu.DM.08G022800) promoters of haplotype 3 (grey), haplotype 4 (orange) and a shortened variant of haplotype 4 (blue), which completely loses its inducibility in heat. Average ± SE is shown. Peak luminescence per well was compared between hap3 and hap4 with a two-sided Wilcoxon rank-sum test, and p-values were FDR-adjusted (Benjamini–Hochberg); control, p.adj = 3.7 × 10⁻¹²; heat, p.adj = 0.0055. ** p.adj < 0.01, *** p.adj < 0.001.

To better understand gene regulation during heat, we performed a motif enrichment analysis of promoters of differentially upregulated alleles. Promoters of 88 strongly (log_2_FC > 2, FDR < 0.05) upregulated alleles vs. 2,294 non-responding alleles (|log_2_FC| < 0.1) were tested for motif enrichment. We identified 31 enriched motifs (Q ≤ 0.05) from 12 transcription factor classes, including binding motifs of HSFs and basic helix-loop-helix factors (e.g., MYC2), both known to be involved in the heat response (Fig. 3b, Supplementary Table S13).

To identify alleles that respond differently to heat, we compared allelic expression between control and heat conditions. Syntelog groups with exactly one copy on at least two haplotypes were tested. 498 (3%) of 15,732 tested pairs showed a differential response to heat (FDR < 0.05 and |Δlog_2_FC| > 1) (Fig. 3c, Supplementary Table S14). This indicates that for most genes all alleles responded similarly to heat, i.e., they were all either not affected, down-, or upregulated.

To confirm the allelic imbalance identified by long-read RNA-seq in heat response, we selected the N²-acetylornithine deacetylase (*NAOD*) gene (Soltu.DM.08G022800) in which haplotype 4 allele was more strongly induced by heat than the other two alleles, while the corresponding allele is absent from haplotype 2 (Fig. 3d). Due to the high variability of the promoters between the two haplotypes, laboratory experiments were required to map the motifs causing the difference. We cloned selected promoters (1,500 bp upstream of the start codon) into vectors upstream of a luciferase reporter, transiently expressed this promoter-reporter fusion in *Nicotiana benthamiana*, and measured luminescence under control and heat conditions. The luciferase assays confirmed the stronger heat response of the haplotype 4 promoter compared with haplotype 3, as predicted by long-read RNA-seq (Fig. 3f). Interestingly, heat inducibility was lost when the haplotype 4 promoter was shortened to 605 bp upstream of the start codon, indicating that the deleted region (605–1,500 bp upstream of the start codon) is required for heat-responsive expression. As this region differs at only a few positions between the haplotype 4 and haplotype 3 promoters, these sequence differences represent strong candidates for the enhanced heat responsiveness of the haplotype 4 allele (Fig. 3e). Consistent with this finding, assay for transposase-accessible chromatin with sequencing (ATAC-seq) revealed a major peak of chromatin accessibility in heat-treated samples for both alleles, located approximately 1,000–1,300 bp upstream of the start codon (Fig. 3e). This region overlaps the deleted segment, further supporting its role in heat induction.

### Long-read RNA-seq resolves the heat response of individual members of multigene families

We next examined individual members of multigene families, which short-read RNA-seq resolves poorly. One such family, the heat shock proteins, is regulated by HSFs, so their promoters are expected to carry HSF binding motifs. To study their regulation, we combined chromatin accessibility data, haplotype-resolved promoter sequences and expression profiles. We observed that the heat shock protein genes (*HSP90*, *HSP70*, *HSP101* and DNAJ-containing proteins) that were significantly upregulated in heat-treated samples all had ATAC-seq coverage colocalized with motifs of HSFs (Fig. 4a). Notably, the motif shared by two HSFs from tomato, SolycHSFB2b and SolycHSFA7, was found in most ATAC-seq coverage peaks close to the TSS and was also found to be enriched in strongly heat-upregulated alleles (Fig. 3b). The alleles of each gene responded similarly to heat (comparable log_2_FC heat/control), the only exception being a DNAJ-containing protein (Soltu.DM.01G039980). Haplotype 2 allele of this DNAJ-containing protein has a 5,536 bp deletion compared to the other haplotypes, potentially affecting heat inducibility (Fig. 4a).

**Fig. 4:**
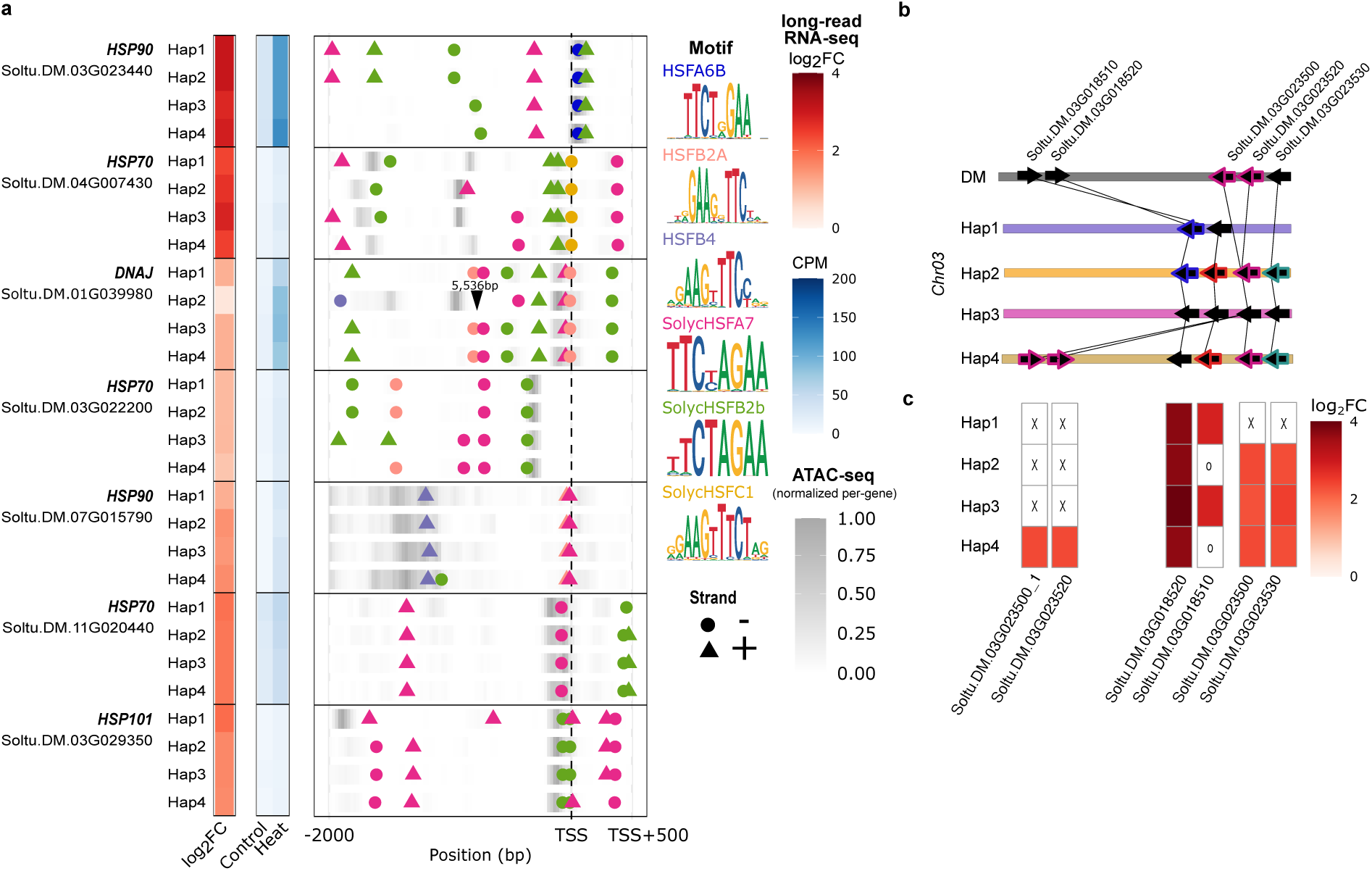
Heat-responsive gene families. **(a)** Log_2_FC, average CPM expression in control and heat, ATAC-seq coverage normalized by gene and six known HSF binding motif matches shown 2,000 bp upstream and 500 bp downstream of the TSS. Motifs identified on the forward (triangle) and reverse (circle) strands of HSFA6B (blue), HSFB2A (red), HSFB4 (purple), SolycHSFA7 (pink), SolycHSFB2b (green) and SolycHSFC1 (yellow). Where motifs overlapped, the one with the highest position weight matrix (PWM) score is shown. For the DNAJ protein (Soltu.DM.01G039980), a 5,536 bp deletion in the promoter of haplotype 2 is marked by a black triangle. **(b)** Haplotype-resolved locations of Kunitz-type protease inhibitor genes on chromosome 3 with differential expression during heat. The same genes are connected by lines and identical alleles share color. Arrow directions correspond to orientation in the genome. **(c)** Heatmap showing log_2_FC and presence/absence (“X”) of Kunitz-type protease inhibitor genes. Lowly expressed genes marked with “0”.

Next, we looked at the Kunitz-type protease inhibitor gene family located on chromosome 3, members of which were differentially expressed during prolonged heat stress. Their copy number varies between haplotypes: haplotype 1 lacks two of the protease inhibitor genes, whereas in haplotype 4 two genes are translocated and duplicated. Haplotypes 2 and 4 share all but one allele, whereas haplotype 3 carries unique alleles (Fig. 4b,c). Among the Kunitz-type protease inhibitors, Cysteine protease inhibitor 8 (*CPI8,* Soltu.DM.03G018520) was the most highly expressed and had the highest log_2_FC during heat across all haplotypes. It is known to be induced by jasmonic acid during biotic stress^26^, and we observed a *MYC2* binding site 714 bp upstream of the TSS within an ATAC-seq peak (Supplementary Fig. S6).

### Prolonged heat stress leads to differential isoform usage for 390 genes

We also explored isoform diversity in potato leaves at the allele level. To identify genes whose major isoform (mRNA splicing variant) differed between haplotypes, we compared expressed isoform structures across haplotypes under control conditions. Of 4,605 genes tested, 154 differed significantly in their major isoform, mostly within the untranslated regions (UTRs) (Fig. 5a, Supplementary Table S15).

**Fig. 5:**
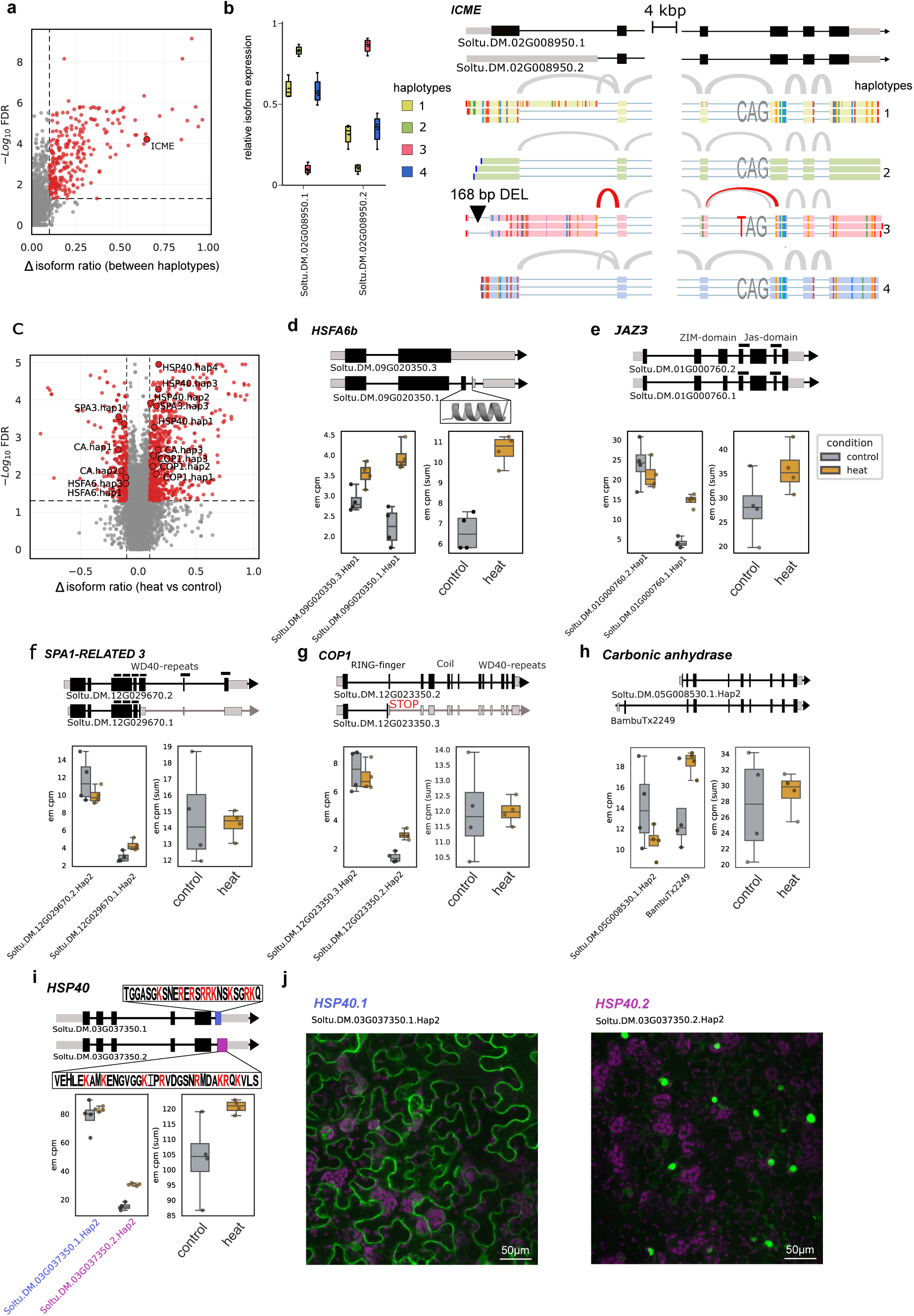
Isoform expression across alleles of different haplotypes and under heat stress. **(a)** Volcano plot of genes with a difference in major isoform expression between alleles under control conditions. The marked gene is further investigated in panel b. **(b)** Left: Relative major isoform expression per allele of *ICME* (Soltu.DM.02G008950). Right: Aligned reads from the four haplotypes and splice sites for the same gene. Splice sites specific to haplotype 3 marked in red. For clarity, 4,000 bp of exonic sequence with identical splicing across all haplotypes were omitted from the visualization. DEL: 168 bp deletion in the 5′ UTR of haplotype 3. **(c)** Volcano plot of differential isoform usage between heat and control conditions. **(d–i)** Individual genes with heat-responsive differential isoform usage, each shown as a gene model with the alternative splice event, isoform CPMs for one haplotype and gene CPMs in control versus heat. Black: CDS, grey: UTRs; **(d)** *HSFA6B* (Soltu.DM.09G020350) where isoform 1 has an additional coding exon; **(e)** *JAZ3* (Soltu.DM.01G000760), with an alternative isoform with a shortened third exon; **(f)** *SPA1-RELATED 3* (*SPA3*; Soltu.DM.12G029670), with a truncated isoform retaining three of seven WD40 repeats; **(g)** *COP1* (Soltu.DM.12G023350), with an alternative 3′ splice site producing a shorter isoform with a premature stop codon; the CDS is shown from the start codon shared with the full-length isoform, the likely translation start, rather than the annotated longest CDS. **(h)** carbonic anhydrase (Soltu.DM.05G008530) with an isoform with alternative TSS; **(i)** *DNAJ/HSP40* chaperone (Soltu.DM.03G037350), with an alternative 3′ splice site introducing a frameshifted, extended C-terminus; Lysines/K and Arginines/R are marked in red. **(j)** Confocal microscopy images of *N. benthamiana* leaves transiently expressing *HSP40* isoform 1 (Soltu.DM.03G037350.1, left) or isoform 2 (Soltu.DM.03G037350.2, right) fused to mTagBFP2. Overlay of the fluorescent protein channel (green) and chlorophyll autofluorescence (purple). All channels are shown in pseudocolor. Separate channels are shown in Supplementary Fig. S9; raw images are deposited in Zenodo.

For example, the gene Soltu.DM.02G008950, coding for an isoprenylcysteine methylesterase (ICME), has an allele-specific isoform 2 predominantly expressed on haplotype 3 differing in two splice sites from the other haplotypes (Fig. 5b), one of these is the alternative 5′ splice site in the first exon. Haplotypes 1 and 4 had low expression of this isoform and haplotype 2 did not express it at all. There is no splice site mutation in the first intron in the second isoform to explain its alternative splicing difference in haplotype 3, but it has a 168 bp deletion in the 5′ UTR located 19 bp upstream of the start codon. In addition, haplotype 3 expressed transcripts with a shortened exon 9 due to a variation in the 3′ splice site of exon 9 (Fig. 5b). Similarly, in Atlantic, haplotype 4 expressed the same alternative isoform (Supplementary Fig. S7).

To identify isoform usage changes induced by heat, we compared isoform usage ratios across all expressed genes, identifying 390 of 13,115 tested genes with differential usage (change > 10%, FDR < 0.05; Fig. 5c, Supplementary Table S16). For each of these genes, isoform usage shifted in the same direction in every expressed haplotype. We examined genes with known heat-response functions or previously reported alternative splicing, as described below. In *HSFA6B* (Soltu.DM.09G020350), the two major isoforms were upregulated under heat, with increased use of the isoform carrying an additional C-terminal exon relative to the control-dominant isoform (Fig. 5d); this splicing replaced the last 9 residues with 23 residues predicted to form an amphipathic α-helix (Supplementary Fig. S8). For the jasmonate-ZIM-domain protein *JAZ3* (Soltu.DM.01G000760), heat increased the abundance of isoform 1 (Soltu.DM.01G000760.1), which uses an alternative 3′ splice site of the third exon that shortens the exon by 96 nt, while expression of isoform 2 decreased; the TIFY and ZIM domain regions are unaffected, and the protein is an ortholog of jasmonate-ZIM-domain protein 3 in Arabidopsis (Fig. 5e). *SPA1* (Suppressor of phyA-105)*-RELATED 3* (Soltu.DM.12G029670), an ortholog of Arabidopsis *SPA3*, showed increased expression of an alternative isoform (Soltu.DM.12G029670.1) in haplotype 2 under heat stress; this isoform uses an alternative 5′ splice site within the fourth intron, potentially introducing a premature stop codon, and the predicted truncated protein retains three of the seven WD40 repeats present in the longer isoform (Fig. 5f). In *COP1* (Constitutive Photomorphogenesis 1; Soltu.DM.12G023350), increased use of an alternative 3′ splice site in the second exon under heat extends this exon from 78 to 114 nt, introducing a premature stop codon in Soltu.DM.12G023350.2 (Fig. 5g). Finally, in beta-carbonic anhydrase (Soltu.DM.05G008530), a novel isoform (BambuTx2249) arising from an alternative transcription initiation site accounted for the majority of transcripts on haplotype 2 under heat, compared with roughly equal isoform proportions under control conditions, and this alternative TSS produces a coding sequence (CDS) with a different N-terminus (Fig. 5h).

### Heat-induced splicing relocalizes a DNAJ/HSP40 chaperone to the nucleus

Heat also changed isoform usage of the DNAJ/HSP40 chaperone Soltu.DM.03G037350. Its two main isoforms differ at the 3′ splice site of the fifth intron: isoform 2 (Soltu.DM.03G037350.2) uses an alternative site 4 nt downstream of the site used by isoform 1 (Soltu.DM.03G037350.1). Under heat, the usage of isoform 2 increased in all four haplotypes (FDR ≤ 0.003), on average from 20% to 35% of transcripts, whereas that of isoform 1 decreased on average from 72% to 60%; isoform 1 nevertheless remained the main isoform in both conditions. The alternative splice site causes a frameshift that gives isoform 2 a longer, different C-terminus (Fig. 5i). This difference was predicted to alter subcellular localization: isoform 1 was predicted to localize about equally to the nucleus and cytoplasm, whereas isoform 2 was predicted to localize mainly to the nucleus (Supplementary Table S17).

To test the hypothesis that the two expressed isoforms lead to different localization of HSP40, we fused the coding sequences of both isoforms to the fluorescent tag mTagBFP2 under the control of the CaMV 35S promoter. The constructs were transiently expressed in *N. benthamiana* leaves, and their subcellular localization was examined using confocal microscopy under control conditions. The two isoforms localized to distinct compartments: isoform 1 to the cytoplasm and isoform 2 to the nucleus (Fig. 5j, Supplementary Fig. S9).

## DISCUSSION

Prolonged, moderate heat is the form of heat stress most relevant to potato cultivation, yet its molecular effects have so far been studied almost exclusively at the whole-gene level^4,27,28^. Zagorščak et al. performed integrative multi-omics profiling of heat, drought and waterlogging in Désirée and identified stress-specific transcriptional, proteomic and metabolic signatures and their hub regulators, but quantified each gene as a single unit, without distinguishing its alleles or isoforms^27^. More recent work has begun to resolve stress responses at the isoform level^10^, yet how the individual alleles of a gene contribute to heat response in potato remained unexplored.

Here, we combined a haplotype-resolved Désirée reference genome with long-read RNA-seq to quantify expression at the allele and isoform level under control conditions and after 7 days of heat (Fig. 1a). Because long reads can be assigned to individual alleles more reliably than short reads, long-read RNA-seq is particularly beneficial for allele-level quantification in polyploids and in diploids with duplicated genes (Fig. 1b)^13,29^. In our data, short reads systematically underestimated allelic ratio differences (Supplementary Note S1), likely because reads that cannot be uniquely assigned dilute allele-specific signal.

Haplotype-resolved reference genomes, combined with long-read RNA-seq, in principle allow promoter variation between alleles to be linked to expression differences. In organisms with little sequence variation between haplotypes, alleles differ at only a few positions, so expression differences between them can be more easily attributed to specific *cis*-regulatory variants^30^. However, in highly heterozygous organisms such as commercial potato cultivars, alleles typically differ at many promoter positions at once (Supplementary Fig. S4a, Supplementary Table S10), and for most genes it is unknown which transcription factors regulate them and where they bind. It is therefore difficult to determine which of the many sequence differences, if any, causes an observed expression difference. Consistent with this, many alleles with promoter SVs were expressed at similar levels (Fig. 2c), showing that an SV alone does not predict allelic imbalance unless it affects functional *cis-*elements.

While *cis*-elements in promoters influence expression level, *cis*-elements (e.g. splice sites) within exons or introns can regulate which isoform is expressed. We identified genes whose major isoform differed between haplotypes under control conditions and, in some cases, linked this to specific sequence differences (Fig. 5a, Supplementary Table S15). In ICME (Soltu.DM.02G008950), the haplotype 3 isoform differed from the other haplotypes at two splice sites (Fig. 5b). One difference likely resulted from a C>T variant at position −3 of the 3′ splice site (CAG>TAG) (Fig. 5b), weakening the splice site and leading to the selection of a cryptic splice site^31^. If this isoform is non-functional, the other three alleles would still provide the canonical isoform, lowering but not abolishing ICME activity. As Arabidopsis ICME (AT5G15860) promotes abscisic acid (ABA) signaling^32^, a lower dosage of potato ICME might subtly dampen ABA responsiveness.

If promoter sequence contributes to allelic imbalance, alleles that share an identical promoter should show similar imbalance in different cultivars. Such shared promoters are expected in potato, because historical European cultivars have low haplotype diversity^33^. Indeed, we identified identical promoters between Atlantic and Désirée, and the relative expression of alleles sharing a promoter was highly correlated between the two cultivars (Fig. 2e, Supplementary Table S11), indicating that allelic imbalance is at least partly encoded in *cis* and is reproducible across cultivars. Genes imbalanced in both cultivars included the RPM1-interacting protein *RIN4* (Soltu.DM.06G033690), where one haplotype was approximately twice as highly expressed (Fig. 2f). Allelic dosage of such a gene may matter for disease resistance, as *AtRIN4* (AT3G25070) regulates both PAMP-triggered and effector-triggered immunity^34,35^.

Interestingly, only 3% (498) of the allele pairs tested showed a differential allelic response to heat (Fig. 3c, Supplementary Table S14), suggesting that the *cis*-regulatory regions of alleles of a gene either share most of their heat-responsive elements or lack such elements altogether. We restricted this analysis to syntelog groups in which alleles differ in sequence and are present as exactly one copy on at least two haplotypes, because only for these can reads be assigned unambiguously to individual alleles. This conservative selection may enrich for conserved genes, so the proportion of divergent allele pairs genome-wide could be higher. In a potato study across a range of abiotic and biotic stresses, alleles of the same gene were assigned to different coexpression network modules, pointing to divergent allelic regulation across conditions^20^. Beyond potato, divergent stress responses between gene copies have so far been described mainly for homoeologs in allopolyploids. In hexaploid wheat, homoeologs from the three subgenomes showed extensive expression bias and expression partitioning during heat and drought acclimation^36^. In the allopolyploid grass *Brachypodium hybridum*, the early transcriptional response to prolonged heat was driven by homoeologs inherited from the heat-tolerant progenitor *B. stacei*^37^. In allotetraploid *Brassica napus*, homoeologs differed in expression under heat, cold and drought, with expression biased towards the ‘A’ subgenome under all conditions^38^. Unlike homoeologs, which derive from the genomes of distinct progenitor species, the haplotypes of autotetraploid potato are alleles of a single genome and are less divergent, which may explain why most allele pairs in Désirée shared their heat response. However, not all haplotypes of cultivated potato are equally divergent, as some stem from introgressions of wild Solanum species, many introduced through breeding for disease resistance^39^. Such wild-derived haplotypes may carry *cis*-regulatory elements that differ from those of the cultivated gene pool and could account for some of the few allele pairs with a divergent heat response.

Identifying alleles that respond differently to environmental stress can reveal favorable variants. Using luciferase assays, we confirmed the differential heat response of one such allele pair in a potato N²-acetylornithine deacetylase homolog most similar to Arabidopsis *NAOD/CGP1* (AT4G17830) (Fig. 3f). AT4G17830 was originally reported to function in ornithine biosynthesis^40^ and was subsequently identified as a cysteinylglycine (Cys-Gly) dipeptidase involved in the cytosolic glutathione (GSH) degradation pathway^41^. The promoter alleles differed in their heat inducibility relative to their basal activity under control conditions. Given that Arabidopsis mutants with reduced glutathione levels show impaired acclimation to high temperatures^42^, haplotype-dependent regulation of this gene could contribute to differences in GSH turnover and redox homeostasis under heat stress in potato. These findings therefore highlight *NAOD* alleles as candidate loci for further investigation of the genetic basis of heat tolerance and their potential utility in breeding heat-resilient cultivars.

Long-read RNA-seq and phased reference genomes also resolve expression changes within gene families, such as the two strongly heat-responsive families examined here: heat shock proteins and Kunitz-type protease inhibitors. The heat shock motif TTCTAGAA, which was also enriched among strongly heat-upregulated alleles (Fig. 3b) and previously identified as an HSF motif in tomato^43^, was present within an ATAC-seq peak in almost all HSP promoters (Fig. 4a). This motif corresponds to a gapless variant of the canonical heat shock element (AGAAnnTTCT), lacking the spacer nucleotides between the two inverted repeats. This suggests that the motif also drives heat-induced expression in potato under prolonged heat.

Kunitz-type protease inhibitors are among the most abundant defense proteins in potato, where they inhibit serine and cysteine proteases and accumulate in response to wounding and pathogen attack^44–46^. Copy number of this family varies between the four Désirée haplotypes, and several haplotypes share identical alleles, yet each gene copy could still be quantified separately (Fig. 4b,c). Several, but not all, members of this family were strongly upregulated under prolonged heat. Cysteine protease inhibitor 8 (*CPI8*, Soltu.DM.03G018520) was the most highly expressed member and showed the strongest heat induction, and we identified a *MYC2* binding site in an accessible region of its promoter (Supplementary Fig. S6), consistent with its reported jasmonate-dependent regulation^26^. Short-read RNA-seq could not resolve these copies, as most of their reads map to multiple alleles.

Alternative splicing is an important mechanism for regulating the heat stress response in plants^9^. For genes with differential isoform usage, all haplotypes responded similarly (Supplementary Table S16). Heat-induced isoform switching was therefore largely shared among alleles, mirroring the uniform allelic response at the expression level. This contrasts with allopolyploid crops, where homoeologs often show partitioned splicing responses to heat, as in wheat^47,48^ and Brassica napus^38^.

In wheat, alternative splicing of the ortholog *TaHSFA6e* under heat stress produces two main transcripts; the longer, carrying an additional 14-aa C-terminal amphipathic helix, shows enhanced transcriptional activity on *HSP70*^49^. We similarly observed a heat-induced increase in an *HSFA6B* isoform with a different C-terminus predicted to form an amphipathic α-helix (Fig. 5d, Supplementary Fig. S8), suggesting that this splicing event may be conserved. In tomato, temperature-dependent splicing of *HSFA2* yields a cytosolic isoform involved in thermomemory and a nuclear isoform driving the early heat response, and intronic variants affecting this splicing co-segregate with thermotolerance^50^. Similarly, the two isoforms of the DNAJ/HSP40 chaperone Soltu.DM.03G037350 were predicted to localize to different subcellular compartments (Supplementary Table S17). Confocal microscopy confirmed this: isoform 1, with a shorter C-terminus, localized to the cytoplasm; isoform 2, induced in heat (Fig. 5i), localized to the nucleus (Fig. 5j, Supplementary Fig. S9). In humans, the same pattern occurs in a DNAJ/HSP40: the longer DNAJB6a isoform is nuclear, whereas DNAJB6b, which lacks an NLS, is cytoplasmic^51^. *DNAJB6a* overexpression also increases nuclear import of another protein^52^. In rice, salt stress induces nuclear import of the chaperone OsDNAJ15, where, together with OsBAG4, it enhances DNA binding of the transcription factor OsMYB106 and thereby promotes salt tolerance^53^. We therefore hypothesize that the nuclear isoform of potato HSP40 may similarly facilitate nuclear import of other proteins or transcription factor binding, and thereby contribute to gene regulation.

## CONCLUSIONS

In summary, long-read RNA-seq mapped to a haplotype-resolved Désirée genome resolves allele– and isoform-level expression that short-read sequencing fails to do accurately. Allele-resolved expression data can therefore help identify haplotypes that confer heat tolerance for potato breeding.

## SUPPLEMENTARY FILES

### Supplementary Tables

Supplementary Table S1: Sequencing and mapping statistics.

Supplementary Table S2: Heat-responsive genes, short-read RNA-seq, DM reference.

Supplementary Table S3: Heat-responsive genes, long-read RNA-seq, DM reference.

Supplementary Table S4: Heat-responsive genes, short-read RNA-seq, haplotype-resolved reference.

Supplementary Table S5: Heat-responsive genes, long-read RNA-seq, haplotype-resolved reference.

Supplementary Table S6: Functional enrichment of heat-responsive genes.

Supplementary Table S7: Synteny classification of genes.

Supplementary Table S8: Allelic imbalance under control conditions.

Supplementary Table S9: Comparison of allelic imbalance between sequencing technologies.

Supplementary Table S10: Promoter structural variants between haplotypes.

Supplementary Table S11: Allelic expression of shared promoters in Désirée and Atlantic.

Supplementary Table S12: Missing haplotypes in Désirée and Atlantic for alleles with identical promoters.

Supplementary Table S13: Motif enrichment in promoters of heat-induced alleles.

Supplementary Table S14: Allele-level heat response.

Supplementary Table S15: Allele-specific major isoforms.

Supplementary Table S16: Heat-induced differential isoform usage.

Supplementary Table S17: Predicted subcellular localization of isoforms.

Supplementary Table S18: gBlock sequences.

### Supplementary Figures

Supplementary Fig. S1: Novel genes and isoforms and corrected gene models identified by long-read RNA-seq.

Supplementary Fig. S2: Correlation of log2FC between long– and short-read RNA-seq for genes grouped by transcript length.

Supplementary Fig. S3: Comparison of long– and short-read allelic quantification.

Supplementary Fig. S4: Promoter divergence of balanced and imbalanced allele pairs.

Supplementary Fig. S5: Transposable element composition of promoter structural variants.

Supplementary Fig. S6: MYC2 binding site within an ATAC-seq peak upstream of a Kunitz-type protease inhibitor gene.

Supplementary Fig. S7: Allele-specific isoform expression of ICME (Soltu.DM.02G008950) in Atlantic.

Supplementary Fig. S8: Predicted protein structures of Désirée HSFA6B isoforms.

Supplementary Fig. S9: Separate channels of the confocal microscopy images in Fig. 5j.

Supplementary Fig. S10: RNA quality and batch-effect correction of long– and short-read RNA-seq.

### Supplementary Notes

Supplementary Note S1: Comparison of allele-specific expression between long– and short-read RNA-seq.

## AUTHOR CONTRIBUTIONS

Conceptualization: N.N., K.G., M.P. Methodology: N.N., V.L., M.P. Software: N.N. Formal analysis: N.N. Investigation: N.N., V.L., T.L., K.S. Resources: T.G. Data curation: N.N., V.L. Visualization: N.N. Writing – original draft: N.N. Writing – review & editing: all authors. Supervision: K.G., M.P. Project administration: K.G., M.P. Funding acquisition: K.G., M.P. All authors read and approved the final manuscript version.

## ACKNOWLEDGEMENTS

The authors wish to thank Marijke Woudsma and Doretta Boomsma from HZPC Research for providing the Désirée plantlets for this study and Klára Panzarová and Lamis Abdelhakim for conducting the experiments at Photon Systems Instruments (PSI) Research Center (Drásov, Czech Republic). We acknowledge the research infrastructures ELIXIR Slovenia and IC Planta (National Institute of Biology) for providing computational resources and plant-growth facilities. The authors would like to acknowledge support of the National Genomics Infrastructure (NGI) for providing assistance in massively parallel sequencing and computational infrastructure. Work performed at NGI has been funded by RFI and Science for Life Laboratory, Sweden.

Moreover, we thank Sophia Sonnewald, Markus Teige and the extended LongTREC network for fruitful discussions.

## METHODS

### Plant material and growth conditions

Potato plants (*Solanum tuberosum* cv. Désirée) were grown as described in Zagorščak et al. (2025). Briefly, in vitro cuttings were planted in 250 ml pots with peat substrate (Klasmann 2) and, after two weeks, transplanted into 3 L pots with peat substrate mixed with sand (3:1). After 32 days of cultivation, plants were assigned to control and heat groups, and leaves were sampled on day 7 of treatment, i.e. 39 days after the in vitro cuttings were planted in substrate. Control plants were cultivated at 22/19 °C (day/night) and heat-treated plants at 30/28 °C (day/night). In both groups, watering was maintained up to 60% field capacity, light intensity was set at 330 µmol m⁻² s⁻¹ PPFD with a 16 h photoperiod, and relative humidity was maintained at 55%. Four biological replicates per condition were collected. For each replicate, the second and third fully developed leaves were harvested and flash-frozen in liquid nitrogen, and leaf tissue was homogenized and aliquoted.

*Nicotiana benthamiana* plants were grown from seeds, and individual seedlings were transferred to pots after two weeks. Throughout their growth, plants were maintained at 22/19 °C (day/night), 55% relative humidity, and an irradiance of 85 µmol m⁻² s⁻¹ PPFD (400–700 nm). For heat treatment, plants were maintained at 28/28 °C (day/night) and 55% relative humidity for 5 days prior to the luciferase assay.

### RNA extraction

Total RNA was extracted and DNase-treated from 80 mg of frozen, homogenized leaf tissue using MagMAX Plant RNA Isolation Kit (Thermo Fisher Scientific) on a KingFisher Apex purification system (Thermo Fisher Scientific). The same RNA samples were previously sequenced with Illumina in 2 × 150 bp paired-end mode^28^. For long-read RNA-seq, full-length cDNA libraries were prepared with the Kinnex full-length RNA kit (PacBio) according to the Iso-Seq protocol and sequenced on three SMRT Cells on a PacBio Revio system at the National Genomics Infrastructure, Sweden, yielding full-length transcript reads.

### Molecular cloning

Genomic DNA (gDNA) was isolated from potato cv. Désirée leaves using the DNeasy Plant Mini Kit (Qiagen) per the manufacturer’s instructions. The *NAOD* (Soltu.DM.08G022800) promoter fragment (1,500 bp upstream of the start codon) was amplified from gDNA by touchdown nested PCR with Phusion High-Fidelity DNA Polymerase (New England Biolabs) using primer pairs Pept-H3-F (5′-caccTACACTGAGGCAAACAAACATCT-3′; forward for haplotype 3) or Pept-H4-F (5′-caccGAACATGTGTAACTACACTGAG-3′; forward for haplotype 4) and Pept-CDS-R (5′-GATAACCAGTGGTCCGCCA-3′; reverse for both haplotypes) followed by PCR using primer pairs Pept-H3-F or Pept-H4-F and Pept-H3-R (5′–GTTAATAATATGGGAATGAAACTAGT-3′; reverse for haplotype 3) or Pept-H4-R (5′-GTTAATAATATGGGAATGAAAGTGC-3′; reverse for haplotype 4). A shorter variant of the haplotype 4 promoter (605 bp upstream of the start codon) was amplified using forward primer 5′-CACCAAAGGGTTAGTTTTTCAAACTGAAC-3′ and reverse primer Pept-H4-R. Coding regions of *HSP40* isoforms 1 and 2 and coding sequence of mTagBFP2 were ordered as gBlocks from IDT (Supplementary Table S18) with overlapping ends and recombined and amplified in a single PCR reaction with primers HSP40-gene-F (5′-CACCATGGTAGACAAAAGCAATGATTA-3′) and mTagBFP2-stop-R (5′-TTAATTAAGCTTGTGCCCCAG-3′). Amplicons were gel-purified using Wizard SV Gel and PCR Clean-Up System (Promega) and cloned into pJET1.2 blunt cloning vector (Thermo Fisher Scientific) according to the manufacturer’s protocol. Chemically competent *E. coli* were transformed with the constructs, plasmids were isolated using the GenElute Plasmid Miniprep Kit (Sigma) and sequenced using the above-listed primers (GATC Biotech). The promoter regions were subsequently amplified from pJET1.2 blunt cloning vector using the above-listed primers and ligated into pENTR/D-TOPO (Thermo Fisher Scientific) following the manufacturer’s protocol. The promoter was recombined from pENTR/D-TOPO into the binary destination vector pGWB435^54^ by LR reaction (Gateway LR Clonase II Enzyme Mix, Thermo Fisher Scientific) following the manufacturer’s protocol. HSP40-mTagBFP2 was recombined from pENTR/D-TOPO into the binary destination vector pK7WG2 (VIB) downstream of CaMV 35S promoter.

### Transient transformation

Homemade electrocompetent *Agrobacterium tumefaciens* LBA4404 cells were transformed with the prepared constructs by electroporation as described before^55^. Overnight cultures of the transformed Agrobacterium were centrifuged for 10 min at 4,000 rpm at room temperature, and the resulting pellets were resuspended in agroinfiltration solution (10 mM MgCl₂, 10 mM MES, and 0.15 mM acetosyringone) to an OD₆₀₀ of 0.3. The prepared Agrobacterium suspensions were incubated with gentle shaking in the dark at room temperature for 2–4 h and subsequently used for agroinfiltration of fully developed leaves of 3–5-week-old *N. benthamiana* plants.

### Luciferase assay

Luciferase assays were performed according to a modified version of the transactivation assay protocol described previously^56^. Briefly, *N. benthamiana* plants agroinfiltrated with Agrobacterium cultures carrying the *NAOD* promoter variants upstream of the luciferase coding sequence were sampled 5 days after heat treatment or control conditions and 4 days after agroinfiltration. Leaf discs were collected using a 0.6-cm-diameter cork borer and placed abaxial side up in white 96-well microtiter plates (Corning) containing MS liquid medium supplemented with 35 µM beetle luciferin (Promega). Luminescence was measured at 30-min intervals using a Centro LB 963 luminometer (Berthold), and the time course of luminescence was plotted for each construct and condition in relative light units normalized to the maximum measured per experiment. For each construct and condition, 24 leaf discs were analyzed. The experiment was performed twice independently. For each construct, the peak luminescence of each of the replicate wells (maximum over the time course) was compared between the haplotype 3 and haplotype 4 promoters with a two-sided Wilcoxon rank-sum test, separately for control and heat, and p-values were adjusted for multiple testing with the Benjamini–Hochberg method^57^.

### Confocal imaging

*N. benthamiana* plants carrying transiently expressed HSP40-mTagBFP2 constructs (isoforms 1 and 2 of *HSP40*) were imaged 3 days after infiltration using the TCS LSI confocal macroscope (Leica) equipped with an ACS APO 20×/0.60 IMM objective. HSP40-mTagBFP2 fluorescence was detected using the 405 nm laser line at 95% power, with an emission window of 440–500 nm and a detector gain of 650 V. Chlorophyll fluorescence was detected using the 488 nm laser line at 90% power, with an emission window of 670–730 nm and a detector gain of 600 V. Images were acquired as Z-stacks encompassing epidermal cells at a voxel size of 5 µm, with the pinhole set to 1 Airy unit, and subsequently presented as maximum-intensity projections. Detailed information on image acquisition is deposited on Zenodo. HSP40 localization was assessed in two leaves from three plants, each set of plants infiltrated with two independent Agrobacterium colonies, and consistent localization patterns were observed.

### Reference genome annotation preprocessing and read mapping

For alignment to a linear reference genome, the reference genome v6.1 of the doubled monoploid potato DM 1-3 516 R44^11^ with the UniTato annotation^23^ was used. Haplotype-resolved Désirée reference genome v1^24^ and haplotype-resolved Atlantic reference genome v3^25^ with gene annotations transferred from UniTato using Liftoff^58^ were used for haplotype-resolved analyses. We used the Liftoff annotation rather than the native Désirée annotation because the native annotation was generated independently for each haplotype, resulting in larger differences in gene models and UTR lengths between alleles; such differences would confound comparisons of allelic expression. Transferring a single annotation to all haplotypes provides consistent gene models across alleles. Liftoff was run separately for each haplotype, providing a chromosome mapping file (-chroms), minimum alignment coverage and sequence identity thresholds of 0.9 (-a 0.9, –s 0.9), and the –copies option to annotate additional gene copies.

Short-read RNA-seq data were processed with the nf-core/rnaseq pipeline v3.27.0^59,60^ (parameters –-pseudo_aligner salmon and –-keepDuplicates) and mapped to the DM and Désirée reference genomes.

Long reads were mapped to the DM reference genome and to the haplotype-resolved Désirée reference genome with the longrnaseq pipeline v1.0.1^61^ run in Nextflow v24.10.6 with the parameter –-technology PacBio. The pipeline performs alignment to the genome and transcriptome with minimap2 v2.29-r1283, novel isoform discovery with bambu v3.4.0^62^, and transcript quantification with oarfish v0.9^63^. For comparisons of allele-specific expression under control conditions, reads were mapped to transcripts with UTRs removed to avoid quantification bias arising from differences in UTR lengths between alleles^13^. ONT long-read cDNA data from cultivar Atlantic were processed in the same way, using –-technology ONT with the corresponding reference genome^25^. Aligned reads were tagged with haplotypes based on primary alignment and visualized with JBrowse 2^64^.

### Batch correction & differential gene expression analysis

Salmon^65^ and oarfish transcript counts were imported into R v4.4.3 using tximport^66^ with transcript-to-gene mappings. Long-read RNA-seq counts aligned to the DM and haplotype-resolved reference genomes were batch-corrected with ComBat_seq from the sva package v3.54.0^67^ using default parameters; the batches used for correction were those visible in RNA concentration (Supplementary Fig. S10a). Batch correction increased the correlation of log_2_FC between long– and short-read RNA-seq (Supplementary Fig. S10b). Short-read RNA-seq counts were not batch-corrected.

For the DM reference genome alignment, genes with fewer than 3 counts per million (CPM) in at least 5 samples were removed prior to analysis. For the haplotype-resolved reference genome, alleles with fewer than 4 CPM in at least 6 samples were removed. Differential expression between heat and control conditions was assessed using the edgeR::voomLmFit function from edgeR v4.4.0^68,69^ with a design matrix modeling the two conditions. An empirical Bayes approach (eBayes) was applied, and differentially expressed genes were defined by an adjusted p-value < 0.05 and |log_2_FC| > 1. Correlation of the edgeR log_2_FC between long-read and short-read samples was calculated with cor(method = “pearson”) for batch-corrected and uncorrected counts.

Functional enrichment of the resulting differentially expressed gene (DEG) sets was performed with clusterProfiler::enricher function^70^ using the UniTato MapMan v4 ontology gene set file (GSEA-compatible, gene level) as the annotation term set, with the background set to all genes tested in the DE analysis. For DEGs using the haplotype-resolved reference, genes that had at least one allele differentially expressed were used. Enriched terms were considered significant at BH-adjusted p-value < 0.05.

### Motif enrichment analysis

Motif enrichment analysis was performed using SEA from the MEME Suite v5.5.9^71^. Promoter sequences were extracted 2,000 bp upstream of the annotated transcription start site (TSS) for two allele sets: heat-induced alleles (log_2_FC > 2, FDR < 0.05) and non-responding alleles (−0.1 < log_2_FC < 0.1). Differential motif enrichment between the two sets was assessed using SEA with the JASPAR CORE (2026) plants database^43^ using default parameters.

### Syntelog identification

Syntelog genes were identified using GENESPACE v1.3.1^72^ within the SyntelogFinder pipeline (v1.0.0)^73^. In brief, genes were first grouped into orthogroups by OrthoFinder2 v3.1.5^74^, and collinear syntenic blocks were identified with MCScanX^75^. Genes were defined as syntelogs when they were both orthologous (assigned to the same orthogroup) and syntenic (located within a shared collinear block); the syntelogs of one gene across the four haplotypes form a syntelog group.

### Allelic expression

Allelic and isoform expression were analyzed with PolyASE v1.3.6^29^; PolyASE functions used for each step are given in parentheses below. For gene-level allele-specific expression analysis under control conditions, syntelog groups were retained (filter_low_expressed_genes) if they had at least 40 raw unique read counts per syntelog group across all replicates under control conditions, were present on all four haplotypes without copy number variation, had a multimapping rate < 0.15 (number of multimapping counts per syntelog group/total counts per syntelog group; calculate_multi_ratios), and had the same transcript length and the same number of transcripts per allele. Allelic ratios were calculated from unique counts (calculate_allelic_ratios, calculate_per_allele_ratios) and tested for imbalance (test_allelic_ratios_within_conditions). Syntelog groups were considered imbalanced if the FDR-corrected p-value was below 0.05 and the ratio difference above 0.1 from the expected 0.25. Short-read RNA-seq counts were processed in the same manner, except that syntelog groups with fewer than 400 salmon em counts were excluded and instead of unique counts, em (expectation maximized) counts were tested for imbalance. For pairwise comparisons (test_allelic_ratios_pairwise), syntelog groups with exactly one copy on at least two haplotypes were tested for imbalance, and alleles with FDR < 0.05 and |log_2_FC| > 1 were considered significantly different.

For cultivar Atlantic, pairwise comparison of allelic expression within leaf control conditions was conducted in the same way as for Désirée for syntelog groups in which all members had more than 40 uniquely mapping reads. For genes tested for pairwise allelic imbalance in both cultivars, promoter sequences 1,500 bp upstream of the TSS were extracted and identical promoters (100% sequence identity and 1,500 bp alignment length) between Atlantic and Désirée were identified by BLASTn v2.12.0^76^. Pearson correlation of log_2_FC was calculated between expression of shared promoter pairs.

### Unique mapping ratios

Unique and ambiguous read counts per transcript were obtained from the oarfish and salmon outputs for one representative sample each, mapped against both the collapsed DM reference and the haplotype-resolved tetraploid Désirée Liftoff annotation. Analysis was restricted to single-isoform genes, and genes with <100 total supporting reads were excluded. For each gene, the unique read fraction was calculated as unique / (unique + ambiguous) reads, and its distribution was compared between long– and short-read RNA-seq, separately for the collapsed and haplotype-resolved references.

### Structural variant identification and promoter analysis

Individual haplotype sequences of the Désirée v1 assembly were aligned pairwise using minimap2 v2.29-r1283 (asm20 preset, –k15 –s50 –z1000 –r3k,100k –O3,20 –-eqx), and SVs were called with SyRI v1.7.0 (-F B)^77^. For this analysis, promoter regions were defined as 2,000 bp upstream of the TSS on the reference as well as query haplotype. SVs were retained if their reference coordinates overlapped the promoter of one gene and their query coordinates overlapped the promoter of a different gene on the same chromosome; cross-chromosomal SVs, SVs lacking query coordinates, and inversions >2,000 bp were excluded. SVs (≥ 50 bp) comprised insertions, deletions, and highly divergent regions (HDRs), i.e., aligned regions with high sequence dissimilarity, as annotated by SyRI. Small variants (< 50 bp; SNPs and indels) were taken from the same SyRI output and assigned to promoter pairs using the same coordinate criteria. Promoter pairs without any called SV, SNP, or indel were considered identical. Differences in the number of SVs and in the distance of SVs to the TSS between balanced and imbalanced promoter pairs were assessed; for promoter pairs without SVs, the number of SNPs and indels and the distance of the first indel to the TSS were compared in the same way. All comparisons were assessed using a one-sided Mann– Whitney U test, with the alternative hypothesis that imbalanced promoter pairs carry more variants than balanced pairs and that their variants lie closer to the TSS.

### Transposable element analysis

For each promoter containing exactly one SV longer than 49 bp, the SV sequence was extracted from SyRI VCF output and queried via BLASTn (e-value ≤ 1×10⁻⁵) against a combined EDTA TE library^24^ built from all four Désirée v1 haplotypes to identify SVs with homology to known transposable elements.

### Differential isoform usage

For isoform-level analysis, genes with a minimum of 100 counts across replicates were retained (filter_low_expressed_genes). Oarfish em counts were used instead of unique counts so that genes with identical sequences on all haplotypes were not excluded. Testing was performed separately for each haplotype: for each isoform, the usage ratio (isoform count / gene total count) was compared between heat and control conditions using a beta-binomial likelihood-ratio test (test_isoform_DIU_between_conditions, which uses betabinom_lr_test from isotools2^78,79^, and p-values were Benjamini–Hochberg corrected. A gene was considered to show differential isoform usage on a given haplotype if at least one of its isoforms had an FDR-corrected p-value below 0.05 and an absolute difference in usage ratio between heat and control conditions greater than 0.1.

### Differential isoform structure between haplotypes

For each syntelog group, the major (or, as fallback, minor) isoform of the most highly expressed haplotype was matched to its structural counterpart on each other haplotype by exon/intron length similarity, and isoform usage (isoform count / haplotype total count) was compared between haplotypes using a beta-binomial likelihood-ratio test (test_differential_isoform_structure). Syntelog groups with FDR < 0.05 and a usage-ratio difference > 0.2 were called significant.

### Pairwise allelic response to heat

For each pair of alleles within a syntelog group, we tested whether their relative expression shifted between control and heat by modeling the ratio of one allele’s count to the pair’s combined count (i / (i + j)) in each condition, using unique long-read counts, with a beta-binomial likelihood-ratio test (test_allelic_ratios_between_conditions, which uses betabinom_lr_test from isotools2^78,79^). Pairs with zero combined coverage in either condition were excluded. P-values were Benjamini–Hochberg corrected across all tested pairs. Δlog_2_FC was defined as the difference between the heat response of the two alleles, Δlog_2_FC = log_2_FC(i) − log_2_FC(j), where log_2_FC is the heat/control fold change of each allele; this is equivalent to the change in the allelic log_2_ ratio (i/j) between heat and control. An allele pair was called as showing a divergent heat response when FDR < 0.05 and |Δlog_2_FC| > 1.

### Prediction of differences between isoforms

Isoform subcellular localization was predicted with DeepLoc 2.1^80^ and functional domains with InterProScan 110.0^81,82^ via their web servers. Structures were compared with AlphaFold Server 2026.08.19^83^. Multiple sequence alignments of promoters and transcripts were done with ClustalW^84^ and DiGAlign^85^.

### ATAC-seq

ATAC-seq samples of control and heat-treated Désirée leaves are publicly available and described in^28^.Paired-end 150 bp reads were processed with the nf-core/atacseq pipeline v2.1.2^59,86^ against the Désirée v1 assembly (scaffolds and organellar sequences removed), retaining multimapping reads with –-keep_multi_map to preserve signal in regions identical between haplotypes; peaks were called with MACS2^87^ and normalized coverage tracks were used for promoter-accessibility analyses.

### Promoter motif scanning and chromatin accessibility profiling

To visualize the relationship between promoter architecture, chromatin accessibility, and heat responsiveness, we selected genes annotated as heat shock proteins (HSPs), chaperones, or heat shock-related transcription factors, retaining those for which at least one haplotype showed a log_2_FC greater than 1 in response to heat. Promoter sequences spanning 2,000 bp upstream and 500 bp downstream of the transcription start site were extracted and scanned for transcription factor binding motifs using position weight matrices derived from JASPAR CORE (2026) plants^43^ entries for SolycHSFA7 (MA2727.1), HSFA6B (MA1664.2), HSFB2A (MA1665.2), SolycHSFC1 (MA2728.1), HSFB4 (MA2020.1), SolycHSFB2b (MA2711.1) and MYC2 (Myc2/la2, MA2736.1). Motif matching was performed with Biostrings^88^ by scanning both the forward and reverse-complement strands at a minimum score threshold of 90% of the maximum PWM score, and the start position, strand, and score of each hit were recorded. In parallel, ATAC-seq signal was extracted from a BigWig track generated from combined heat-sample reads, summarized in 25 bp bins, and normalized per promoter. Motif positions and ATAC-seq signal were then displayed together with allele-specific log_2_FC for the selected genes and haplotypes.

## DATA AVAILABILITY

All sequencing data of Désirée leaves under control and prolonged heat conditions are available in the ENA under BioProject PRJEB127277. The long-read Kinnex RNA-seq data were generated in this study (run accessions ERR18072167–ERR18072174). Short-read RNA-seq data and ATAC-seq data of the same conditions, generated previously^28^ are available under run accessions ERR18076900–ERR18076907 and ERR18072205– ERR18072212.

Publicly available ONT long-read RNA-seq data of Atlantic were obtained from the SRA (SRR14993894, SRR14993895, SRR14996168).

Subsamples of the long-read RNA-seq, the GFF containing novel isoforms and bigwigs of ATAC-seq have been added to the Désirée genome browser (https://desiree.nib.si). The updated annotation (GFF) of the haplotype-resolved Désirée genome, including novel genes and isoforms identified by long-read RNA-seq, and the raw confocal microscopy images of HSP40-mTagBFP2 localization in *N. benthamiana*, together with the full image acquisition settings, are available on Zenodo <u>10.5281/zenodo.23594697</u>.

## CODE AVAILABILITY

Analysis scripts are available on GitHub: https://github.com/NIB-SI/LR-DESIREE-reproducibility.

