## Supplementary Figure S1-S10 for "Haplotype-resolved long-read transcriptomics uncovers allele-specific heat stress induction and a nucleus-targeted chaperone isoform in potato"

### Supplementary Figures

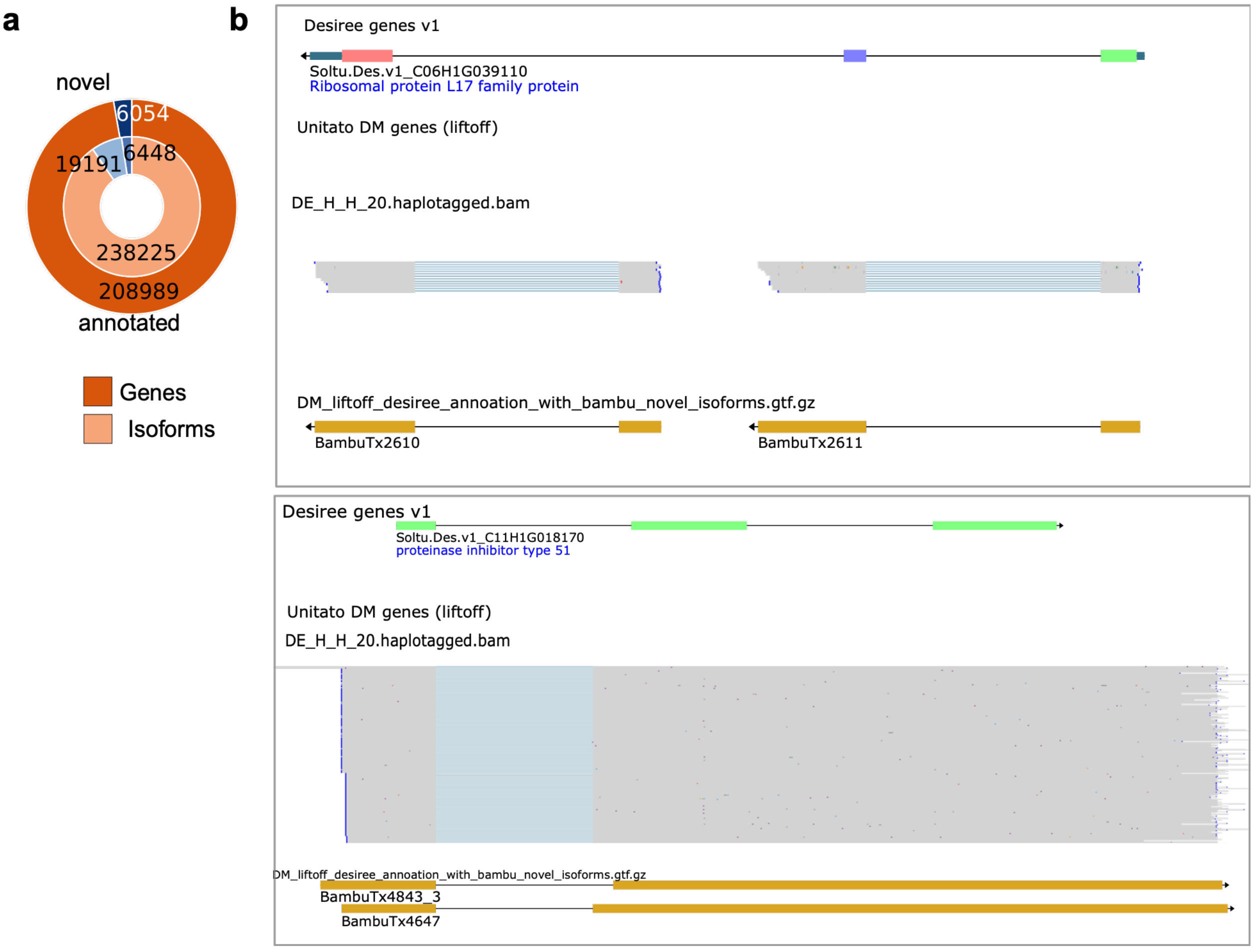


**Supplementary Fig. S1: Novel genes and isoforms and corrected gene models identified by long-read RNA-seq. (a)** Pie chart of annotated and novel genes and isoforms. Dark/light orange: previously annotated genes/transcripts; dark/light blue: novel genes/transcripts discovered by Bambu from long-read RNA-seq and absent from the Liftoff DM annotation. **(b)** Two examples of gene models corrected by long-read RNA-seq relative to incorrectly annotated structures in the Désirée v1 annotation. Top: BambuTx2610 and BambuTx2611 split Soltu.Des.v1_C06H1G039110, suggesting that two adjacent genes were falsely merged into a single model. Bottom: BambuTx4647 and BambuTx4843_3 showing that the gene model Soltu.Des.v1_C11H1G018170 splicing is predicted incorrectly.

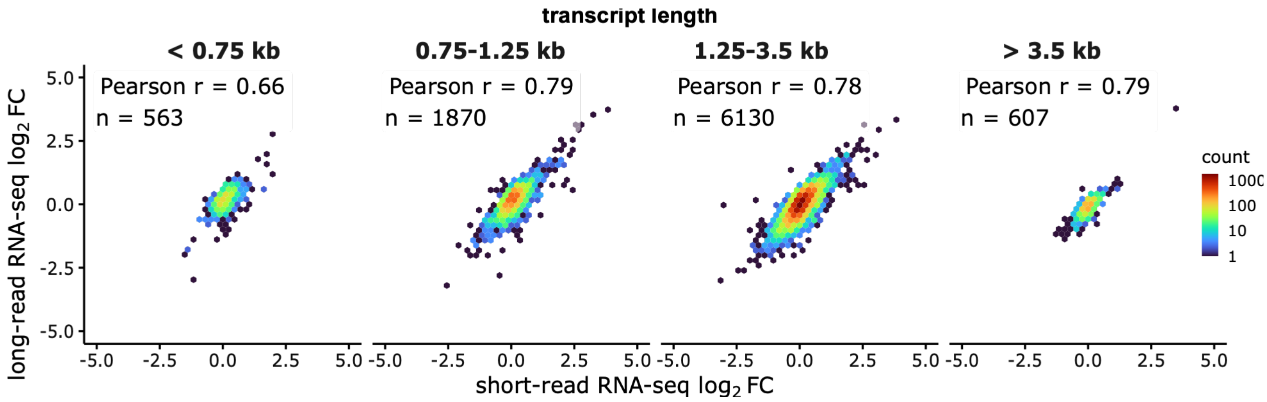


**Supplementary Fig. S2: Correlation of $log_2$FC between long- and short-read RNA-seq for genes grouped by transcript length**. $Log_2$ fold changes ($log_2$FC) were calculated from long- and short-read RNA-seq reads aligned to the DM reference genome.

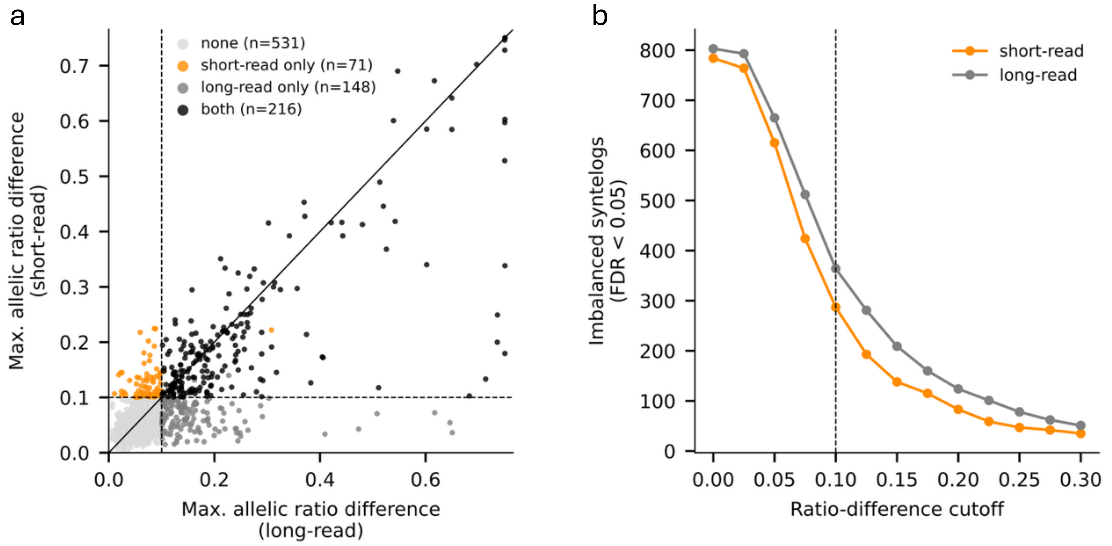


**Supplementary Fig. S3: Comparison of long- and short-read allelic quantification. (a)** Largest allelic ratio difference per syntelog, long- vs. short-read RNA-seq, colored by the method(s) that detected significant allelic imbalance. **(b)** Number of syntelogs with significant allelic imbalance at increasing ratio-difference cutoffs. Dashed lines mark the cutoff used in this study. Ratio difference is an allele's absolute deviation from balanced expression.

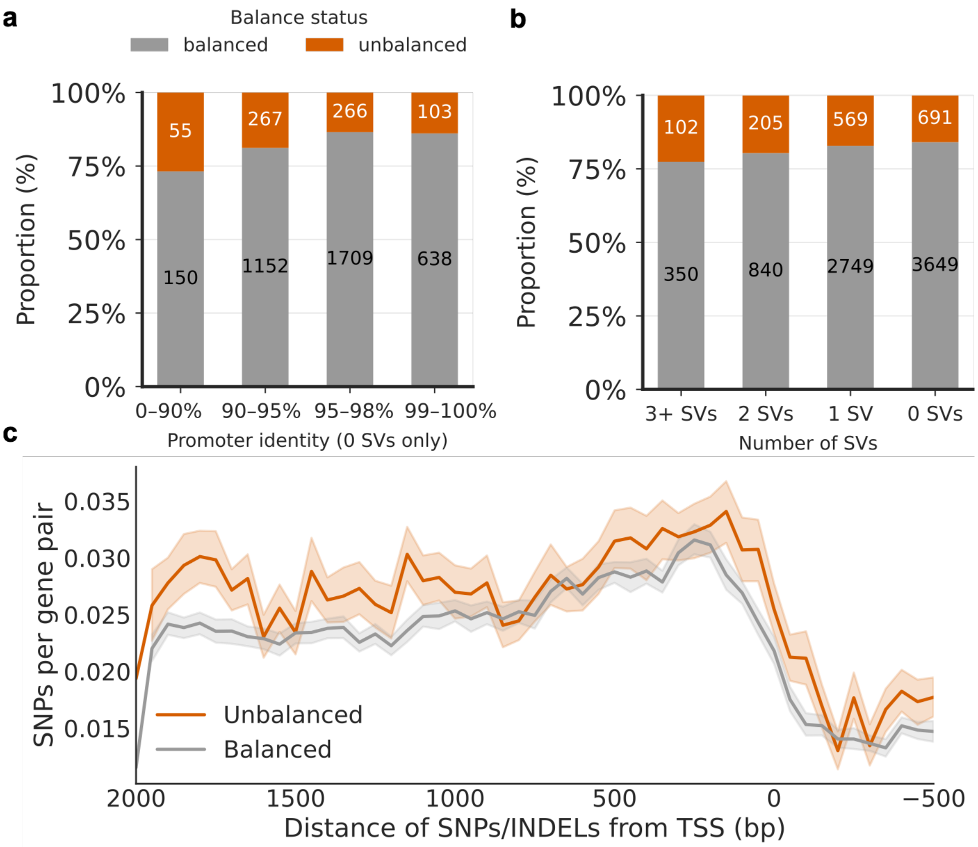


**Supplementary Fig. S4: Promoter divergence of balanced and imbalanced allele pairs. (a)** Number of balanced and imbalanced syntelogs, binned by promoter-pair sequence identity (promoter pairs without SVs only). **(b)** Proportion of balanced and imbalanced syntelogs by number of structural variants (SVs) per promoter pair. **(c)** SNP/INDEL density per gene pair across promoter sequence position relative to the transcription start site (TSS), comparing syntelog pairs with balanced versus imbalanced allelic expression.

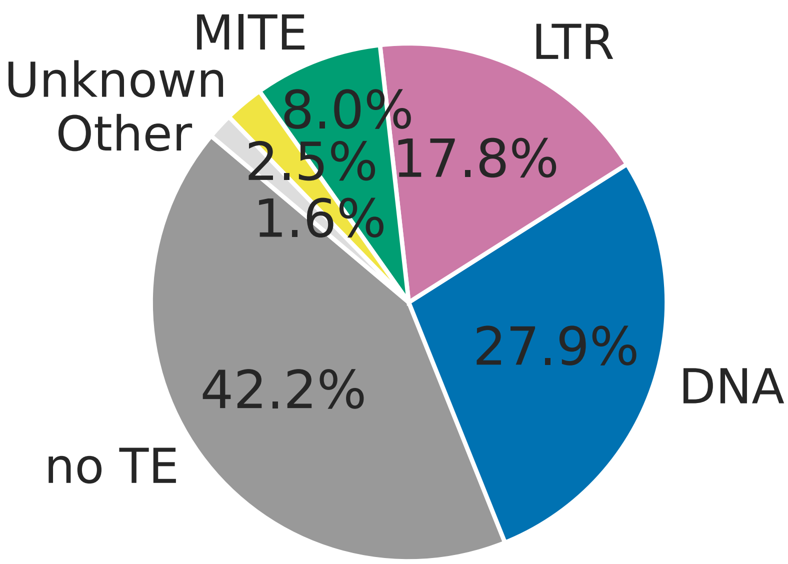


**Supplementary Fig. S5: Transposable element composition of promoter structural variants.** Composition of transposable element classes among structural variants (SVs) identified in syntelog promoter pairs with exactly one SV. "no TE" denotes SVs without transposable element sequence. MITE, miniature inverted-repeat transposable element; LTR, long terminal repeat retrotransposon; DNA, DNA transposon (excluding MITEs). "Other" comprises less abundant TE classes ([e.g. LINE, SINE, Helitron]); "Unknown" denotes TE sequences that could not be assigned to a class.

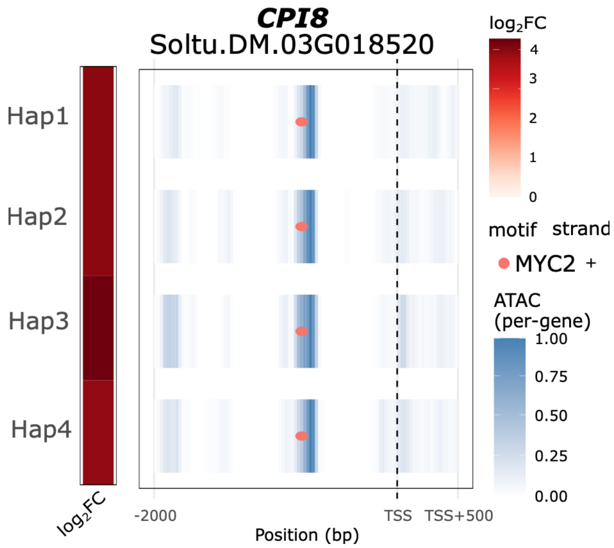


**Supplementary Fig. S6: MYC2 binding site within an ATAC-seq peak upstream of a Kunitz-type protease inhibitor gene.** MYC2 binding motif within an ATAC-seq peak upstream of the Kunitz-type protease inhibitor locus (CPI8, Soltu.DM.03G018520) on chromosome 3.

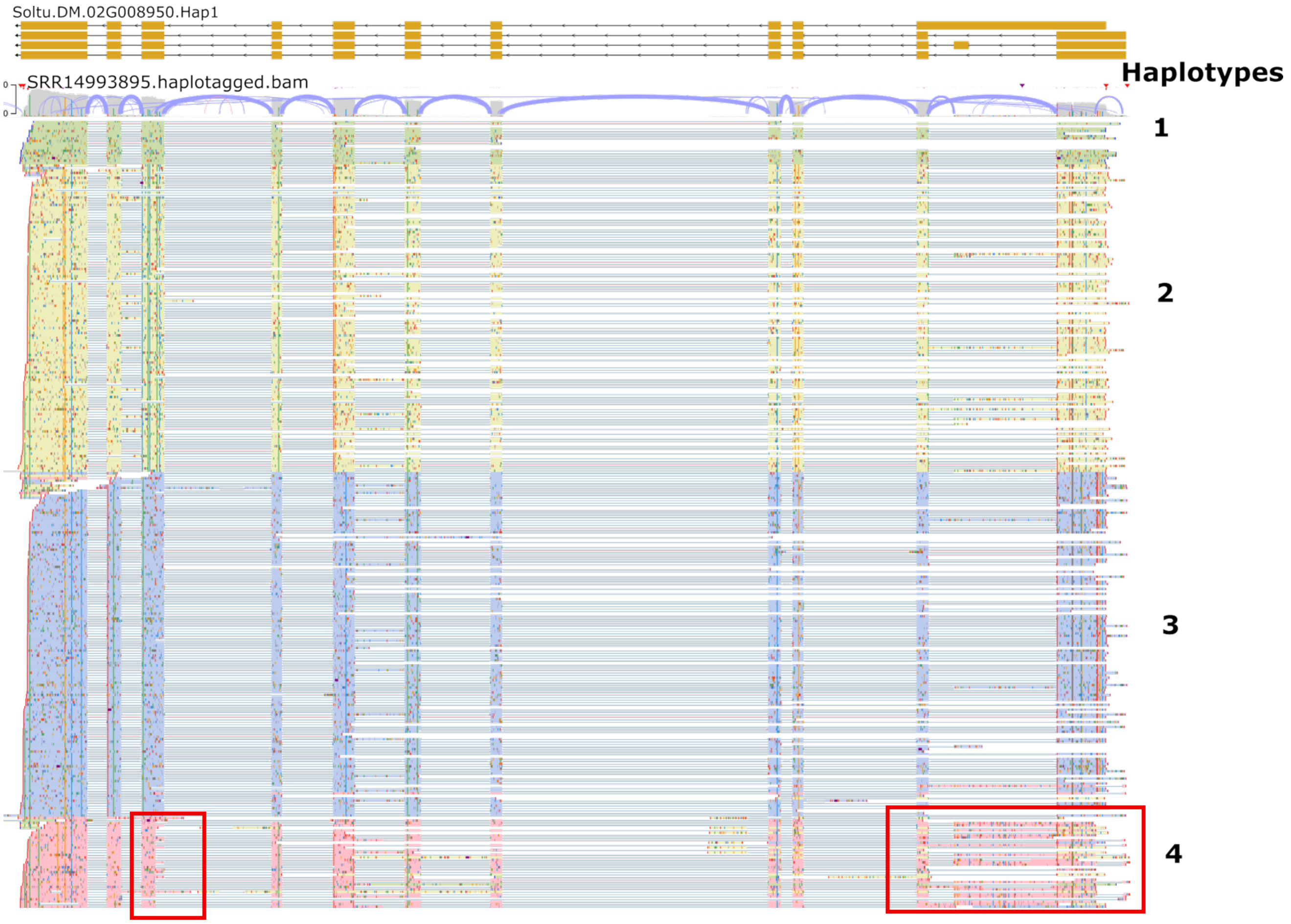


**Supplementary Fig. S7: Allele-specific isoform expression of ICME (Soltu.DM.02G008950) in Atlantic.** JBrowse view of Atlantic long reads aligned to the haplotype 1 allele of Soltu.DM.02G008950. Reads are sorted and colored by haplotype of origin. Atlantic haplotype 4 expresses the same splice variant as Désirée haplotype 3 (Fig. 5b), which differs from the isoforms of the other haplotypes at two splice sites (marked in red).

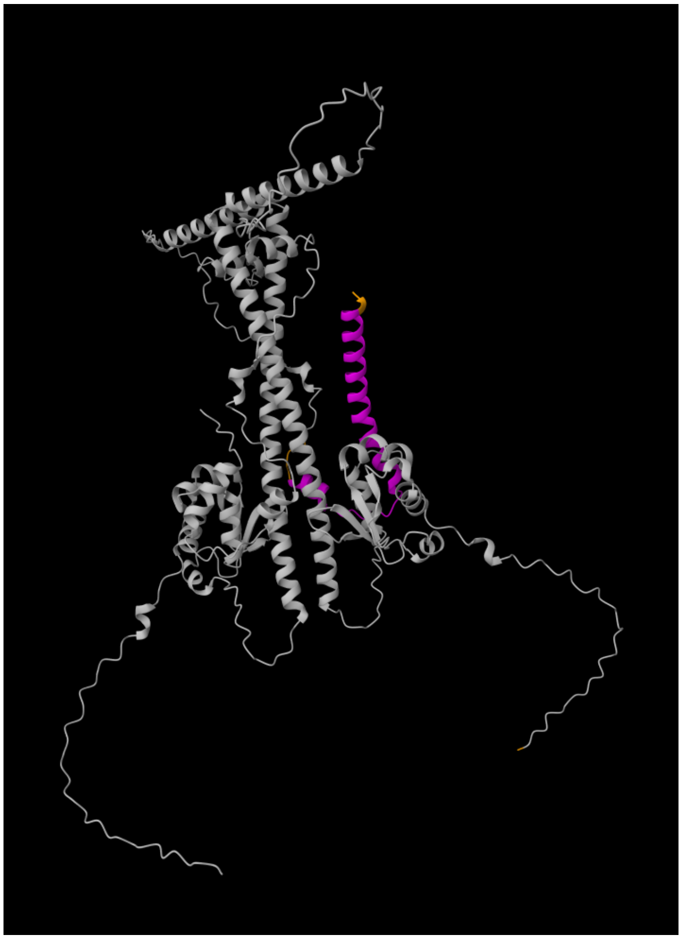

**Supplementary Fig. S8: Predicted protein structures of Désirée *HSFA6B* isoforms.** AlphaFold-predicted structure of a heterodimer formed by two *HSFA6B* (Soltu.DM.09G020350) isoforms. The predicted α-helix of the isoform carrying the additional C-terminal exon is shown in purple.

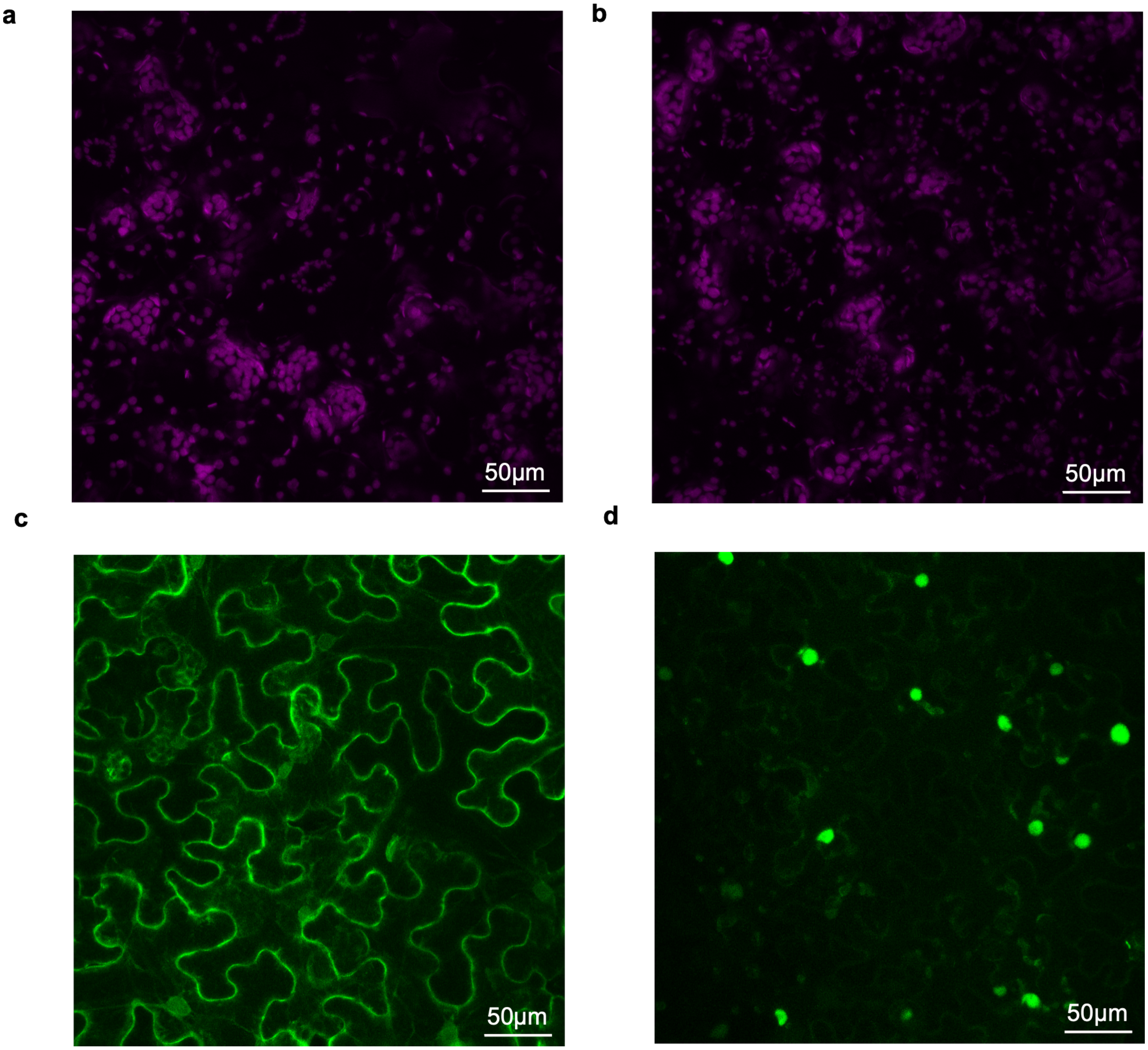


**Supplementary Fig. S9: Separate channels of the confocal microscopy images in Fig. 5j. Individual channels of the confocal microscopy images shown in Fig. 5j** in the main text, from *Nicotiana benthamiana* leaves transiently expressing *HSP40* isoforms fused to mTagBFP2. **(a)** Chlorophyll autofluorescence (purple) of leaves expressing isoform 1 (Soltu.DM.03G037350.1). **(b)** Chlorophyll autofluorescence (purple) of leaves expressing isoform 2 (Soltu.DM.03G037350.2). **(c)** mTagBFP2 fluorescence (green) of isoform 1. **(d)** mTagBFP2 fluorescence (green) of isoform 2. All channels are shown in pseudocolor. Isoform 1 localizes to the cytoplasm and cell periphery, whereas isoform 2 accumulates in nuclei.

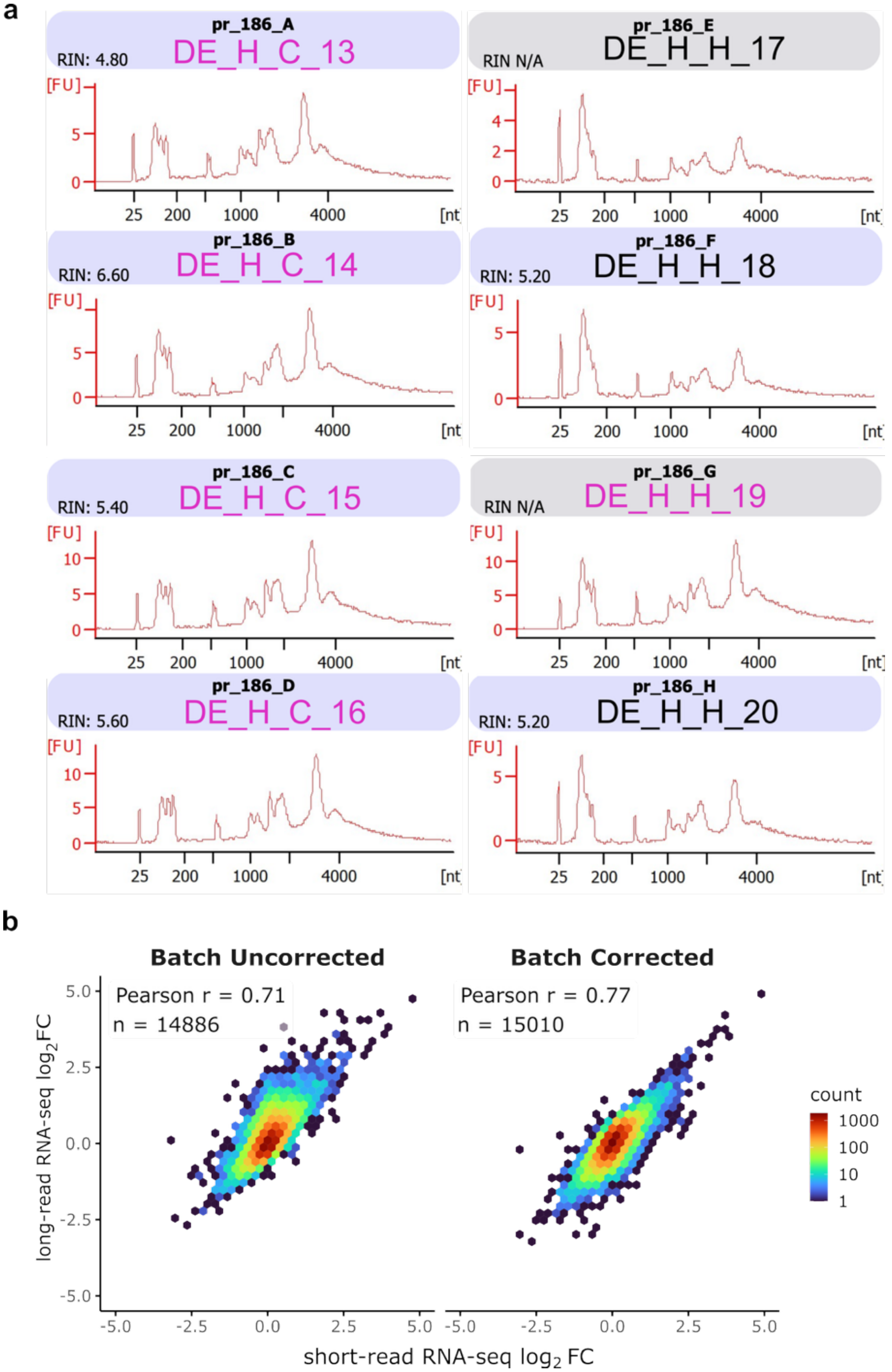


**Supplementary Fig. S10: RNA quality and batch-effect correction of long- and short-read RNA-seq. (a)** RNA integrity number (RIN) of each sequencing sample. Sample names are colored by RNA concentration batch (pink: batch 1; black: batch 2). **(b)** Pearson correlation of $\log_2$ fold changes ($\log_2$FC) between long- and short-read RNA-seq before and after batch-effect correction.
