## Supplementary Note for "Haplotype-resolved long-read transcriptomics uncovers allele-specific heat stress induction and a nucleus-targeted chaperone isoform in potato"

### Supplementary notes

Supplementary Note S1

We compared short- with long-read RNA-seq in allele-specific expression analysis. In the short-read RNA-seq dataset, 5,239 syntelog groups were tested and 861 showed imbalanced expression (FDR < 0.05, no ratio-difference cutoff) (Supplementary Table S9). In total, 966 syntelogs were tested in both long- and short-read RNA-seq. Comparison of allele-specific expression results between short- and long-read RNA-seq showed that the majority of syntelogs with balanced as well as imbalanced expression were concordant between technologies. However, long-read RNA-seq identified a greater number of syntelogs with imbalanced allelic expression than short-read RNA-seq (Supplementary Fig. S3, Supplementary Table S9). Inspection of genes detected as imbalanced exclusively by long-read RNA-seq revealed that, in cases where haplotype transcripts were highly similar and differed by only a small number of SNVs, short-read quantification produced smaller allele ratio differences than long-read quantification. Comparing the largest allele ratio difference per syntelog between technologies showed that 77% of syntelogs with significant imbalance in long-read RNA-seq had a smaller ratio difference in short-read RNA-seq (Supplementary Fig. S3a). Consequently, the number of imbalanced syntelogs (FDR < 0.05) depended on the ratio-difference cutoff: without a cutoff, both technologies detected similar numbers (784 short-read vs. 803 long-read), whereas at a cutoff of 0.1 long-read RNA-seq detected more (287 short-read vs. 364 long-read; Supplementary Fig. S3b). This is consistent with the higher unique mapping rates observed for long-read RNA-seq at the haplotype-resolved reference genome (Fig. 1b), suggesting that longer reads enable more accurate assignment to individual haplotypes, particularly at loci with low sequence divergence.
